# Integrative genetic and epigenetic analysis uncovers regulatory mechanisms of autoimmune disease

**DOI:** 10.1101/054361

**Authors:** Parisa Shooshtari, Hailieng Huang, Chris Cotsapas

## Abstract

Genome-wide association studies in autoimmune and inflammatory diseases (AID) have uncovered hundreds of loci mediating risk^1,2^. These associations are preferentially located in non-coding DNA regions^3,4^ and in particular to tissue-specific Dnase I hypersensitivity sites (DHS)^5,6^. Whilst these analyses clearly demonstrate the overall enrichment of disease risk alleles on gene regulatory regions, they are not designed to identify individual regulatory regions mediating risk or the genes under their control, and thus uncover the specific molecular events driving disease risk. To do so we have departed from standard practice by identifying regulatory regions which replicate across samples, and connect them to the genes they control through robust re-analysis of public data. We find substantial evidence of regulatory potential in 132/301 (44%) risk loci across nine autoimmune and inflammatory diseases, and are able to prioritize a single gene in 104/132 (79%) of these. Thus, we are able to generate testable mechanistic hypotheses of the molecular changes that drive disease risk.

---

The autoimmune and inflammatory diseases (AID) are a group of more than 80 common, complex diseases driven by systemic or tissue-specific immunological attack. This pathology is driven by loss of tolerance to self-antigens or chronic inflammatory episodes leading to long-term organ and tissue damage. Risk variants identified by genome-wide association studies (GWAS) are preferentially located in non-coding regions with tissue-specific chromatin accessibility^7,3,8,9^ and in transcriptional enhancer regions active after T cell stimulation^4^. Formal analyses partitioning the heritability of disease risk across different genomic regions support this enrichment^6^, with excess heritability localizing to tissue-specific DNase I hypersensitive sites (DHS)^5^. Cumulatively, these results suggest that AID pathology is mediated by changes to gene regulation in specific cell populations, but are not designed to identify individual regulatory regions mediating risk or the genes under their control. Several fine-mapping efforts have jointly considered genetic association and epigenetic modification data as a way to identify causal variants ^10,11,12^. However, these efforts use epigenetic mark information to assess whether associated variants are likely to be causal, rather then to identify the regulatory sequences that mediate risk, and the genes they affect.

We have therefore developed a systematic approach to identify regulatory regions mediating disease risk, and thus generate testable mechanistic hypotheses of the molecular changes that drive disease risk (Supplementary Figure 1). For each association, we first calculate posterior probabilities of association from GWAS data and thence the set of markers forming the 99% credible interval (CI)^13,32,14^. We then overlap CI SNPs with DHS in the region to identify which regulatory regions may harbor risk, and from these SNPs calculate the fraction of posterior probability attributable to each DHS. We chose DHS as they are general markers of chromatin accessibility and typically only 150-250 base pairs long, compared to other histone modifications which can span tens to hundreds of kilobasepairs. Next, we identify genes controlled by each DHS by correlating chromatin accessibility state to expression levels of nearby genes. We use the atlas of tissues available at Roadmap Epigenomics Project (REP) data^15,16^, where both DHS and gene expression have been measured in the same samples. Finally, we combine the posterior probability of disease association of each DHS and the correlation between that DHS and the expression levels of nearby genes to calculate a per-gene posterior probability of disease association. This allows us to estimate the probability that a gene mediates disease risk, and to rank genes in a locus by these values.

DHS peaks, as all epigenetic marks, are called in each sample separately^17^. We therefore clustered DHS peaks to identify those corresponding to the same underlying regulatory site, so we could correlate accessibility state of the same site to gene expression data (Supplementary Figure 2). In 56 REP tissues with at least two replicate DHS sequencing runs, we called 22,060,505 narrow–sense 150bp peaks at a false discovery rate FDR < 1%, which fell into 1,994,675 DHS clusters of 250-400bp each, covering 14.8% of the autosomal genome. Of these, 1,079,138 (54.1%) covering 8.5% of the genome passed nominal significance in a statistical replication test (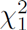 test, *p* < 0.05). This subset explains essentially all the heritability attributable to all peaks in multiple sclerosis and inflammatory bowel disease GWAS (Supplementary Figure 3), indicating they represent the majority of regulatory regions relevant to AID risk. Of these 56 REP tissues, 22 also have gene expression measurements, from which we calculated the correlation between DHS accessibility state and transcript levels. We therefore restricted our present analysis to 1,079,138 DHS clusters and 13771 genes across these 22 REP tissues, though we note our framework can be used with any regulatory feature and expression dataset, and is publicly available.

With this framework, we dissected 301 genome-wide significant associations to one of nine AID, using publicly available summary association statistics from samples genotyped on the Immunochip, a targeted genotyping array^18,19^ (available at immunobase.org; Table 1). We first collated all reported genome-wide significant associations reported for each disease, then restricted our analysis to the loci genotyped at high density on the Immunochip^13,32^. We excluded the Major Histocompatibility Locus, where complex LD patterns make credible interval mapping challenging^20^. For each association, we calculated posterior probabilities of association for all markers and defined credible interval SNP sets^13,14^. We find a median of 4 (standard deviation, sd = 7.8) DHS clusters overlap CI SNPs, out of a median 822 (sd = 205.2) DHS clusters in each 2Mb window around an association (Figure 1A), indicating this data integration step alone vastly reduces the number of potentially disease-relevant regulatory regions.

**Table 1:** regulatory fine mapping resolves 132/301 loci to DHS clusters and 104/132 to single genes across nine autoimmune and inflammatory diseases. We find a substantial proportion of the posterior probability of disease association localizes to DHS clusters in 132/301 (44%) loci. In 104/132 (79%) of these we can prioritize a single gene controlled by the risk-mediating DHS.

| Disease | Dense Immunochip loci |  |  | Regulatory fine-mapping |  |  |
| --- | --- | --- | --- | --- | --- | --- |
|  | Genome-wide significant | >1 credible SNP in a DHS cluster | Posterior on DHS > 0.25 | DHS (genes) prioritized | Posterior on DHS > 0.5 | DHS (genes) prioritized |
| Autoimmune thyroid disease | 8 | 6 | 2 | 6 (5) | 1 | 3 (2) |
| Celiac disease | 31 | 28 | 14 | 68 (42) | 3 | 13 (9) |
| Inflammatory bowel disease | 125 | 97 | 48 | 263 (153) | 24 | 101 (78) |
| Juvenile idiopathic arthritis | 22 | 17 | 7 | 110 (19) | 2 | 19 (6) |
| Multiple sclerosis | 54 | 48 | 23 | 177 (68) | 8 | 56 (24) |
| Primary biliary sclerosis | 15 | 12 | 1 | 3 (4) | 0 | 0 (0) |
| Psoriasis | 24 | 19 | 9 | 64 (29) | 2 | 5 (6) |
| Rheumatoid arthritis | 47 | 40 | 16 | 193 (43) | 6 | 21 (17) |
| Type 1 diabetes | 45 | 34 | 12 | 62 (34) | 7 | 29 (19) |

**Figure 1:**
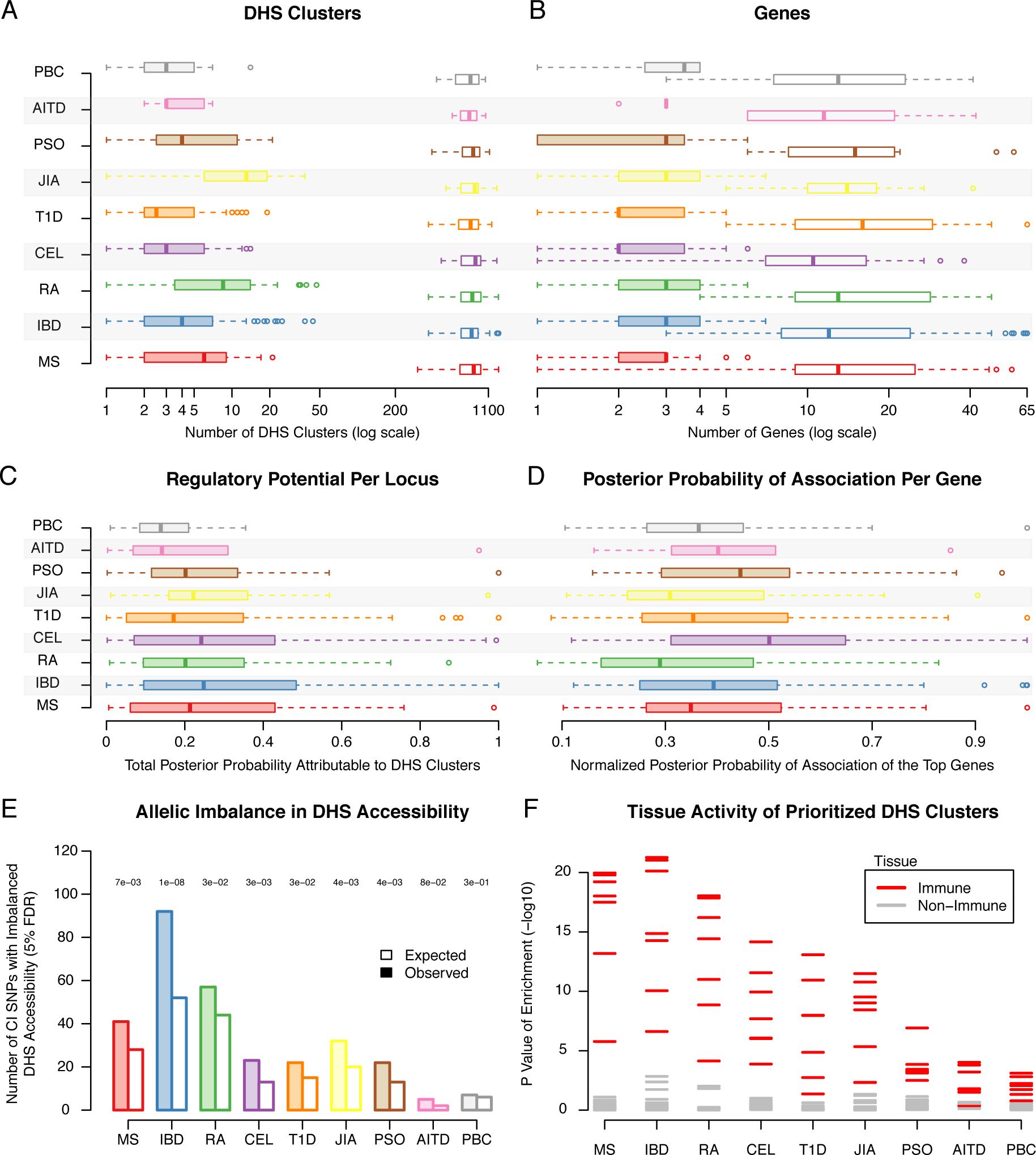
evidence that disease risk is driven by perturbation to specific gene regulatory regions in 301 loci across nine autoimmune diseases. By integrating disease association and DNse I hypersensitivity site (DHS) clusters in a posterior probability framework, we find that. 132/301 (44%) of loci have > 25%) probability of risk being mediated by variants on specific regulatory sequences (53/301, 18%) have >50%) probability). Our framework is able to substantially resolve many autoimmune and inflammatory disease loci from (A) a median of 822 DHS clusters to 4; and (B) from a median of 14 genes to 3. (C) A substantial fraction of the total posterior probability of association in each locus (regulatory potential) localizes to DHS clusters; 132/301 (44%) have >25%) regulatory potential, making them reasonable candidate loci for regulatory effects. (D) This regulatory potential can be attributed to limited number of genes in each locus. The genes with the most regulatory potential, shown here, often account for the majority of signal in the locus making them the only plausible candidate for disease causality in that, region. (E) Credible interval variants that, localize to DHS clusters are more likely to alter DHS accessibility than expected by chance, indicating a functional role. (F) DHS clusters with high regulatory potential are preferentially accessible in immune cell types.

To assess how likely each association is to be mediated by a variants on a regulatory region, we compute their regulatory potential *ρ*, as the proportion of the posterior probability of association localizing to DHS clusters. Consistent with previous observations^3,21,4^ we find that risk often localizes to DHS clusters: over 25% of the posterior is located on DHS clusters in 132/301 (44%) of loci, and over 50% of the posterior in 53/301 loci (18%) (Figure 1C). We reasoned that if DHS clusters harboring CI SNPs actually mediate risk, their accessibility state should be perturbed by the variants they harbor, and they should be accessible in disease-relevant cell populations. We find that CI SNPs on DHS clusters are more likely to induce allele-specific accessibility^22^ (Fisher exact test *p* = 7*e*^−6^, Figure 1E), and that these DHS clusters are more likely to be accessible in immune cell subpopulations (Figure 1F). These results show our approach identifies disease-relevant regulatory events, and support the view that common genetic variants influence disease risk by altering the accessibility of gene regulatory regions.

Having validated that our analysis was identifying genuine regulatory risk effects, we next turned to identifying specific disease-mediating DHS clusters and the genes they control (Supplementary Table 1 and Supplementary Table 2). We focused on the 132 loci where regulatory potential *ρ* > 0.25, as these associations may be mediated by genetic perturbation of a regulatory region. We found that an average of two DHS clusters (*sd* = 4.0) account for > 90% of the total association posterior attributable to all DHS clusters in these loci, indicating we can resolve most loci to a small number of candidate regulators (Supplementary Figure 4). By correlating the accessibility state (open or closed) of each DHS cluster to the expression of nearby genes across 22 REP tissues, we were able to prioritize a median of 3/14 genes per locus (Figure 1B and Supplementary Table 2), and could attribute a gene-wise proportion of posterior probability *γ* > 0.25 to a single gene in 104/132 (79%) loci. Surprisingly, the top-scoring genes are not the closest to the most associated variant in 92/104 (88%) of these cases, suggesting that risk-relevant regulatory regions exerting influence over genes at considerable distances (Table 2). The DHS clusters with high *ρ* values are more likely to be marked as active enhancers of transcription, which can bind distant promoters through long-range DNA looping events^23,24,25^, further supporting this conclusion (Supplementary Figure 5).

**Table 2:**
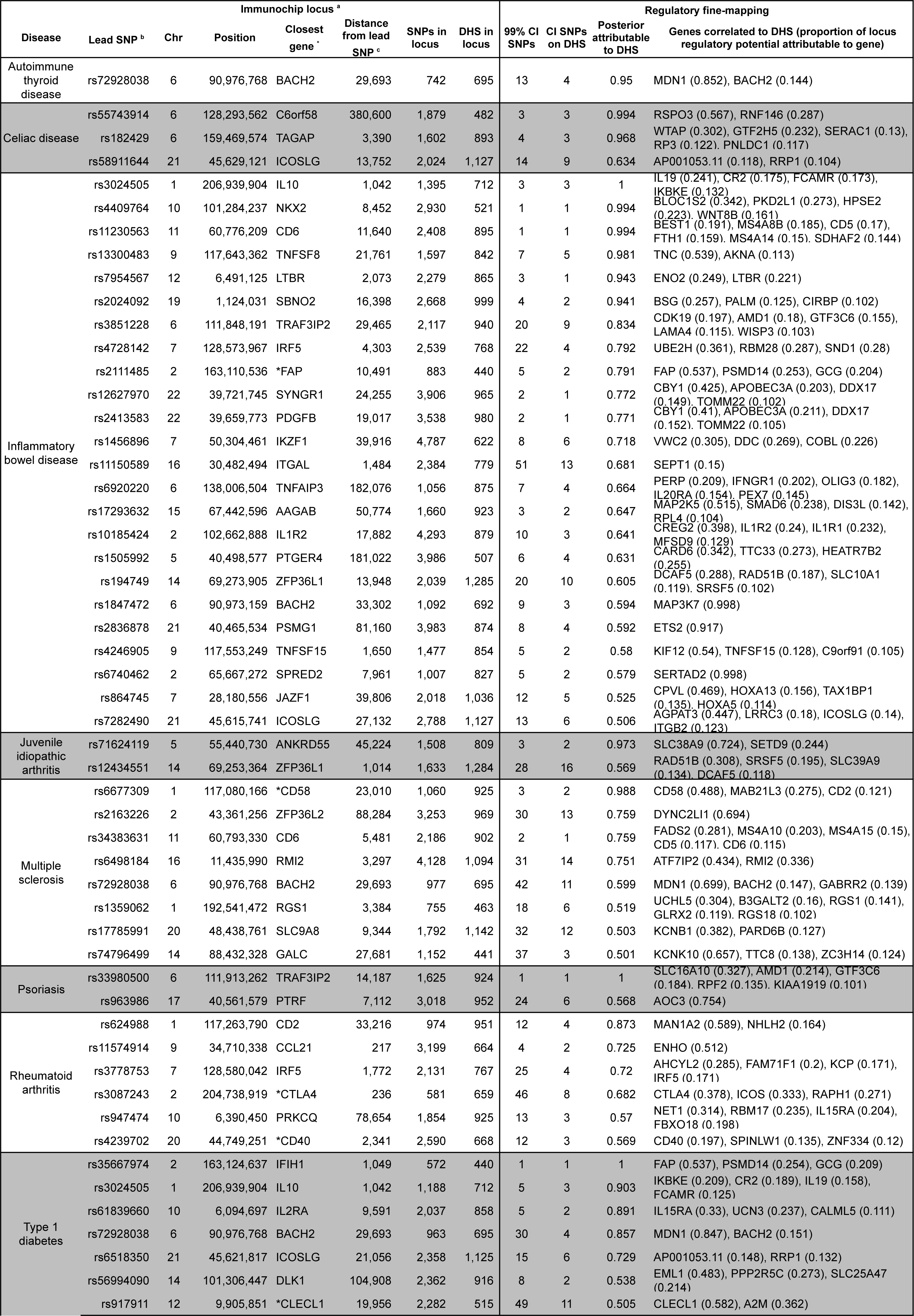
53 risk loci with strong regulatory potential (ρ>0.5) in nine autoimmune diseases. We find that >50% of the posterior probability of association is attributed to SNPs in DHS clusters in 53/301 risk loci. In 19/53, we can attribute the majority (> 50%) of the locus regulatory potential to a single gene. CI: credible interval13, CHR: chromosome; DHS: DNase I hypersensitivity cluster. (a) We define loci as a 2Mb window around the lead SNP, which are (b) associated to disease risk at genome-wide significance in a large meta-analysis, overlap a densely genotyped region of the Immunochip, and are not in the Major Histocompatibility Locus. (c) distance measured between lead SNP and closest gene boundary. Stars indicate genes to which we attribute the highest posterior probability of association. Results for all loci are shown in Supplementary Table 2.

In several cases, we found evidence supporting a previous hypothesis for a causal gene in a locus. For example, we were able to resolve an association to multiple sclerosis (MS) risk on chromosome 1 to two DHS clusters, both of which implicate the *CD58* gene (*γ* = 0.49, Figure 2). *CD58* encodes lymophocyte-function associated antigen 3 (LFA3), a co-stimulatory molecule expressed by antigen presenting cells, mediating their interaction with circulating T cells by binding lymophocyte-function associated antigen 2 (LFA2)^26^. The latter is encoded by the *CD2* gene immediately proximal to *CD58*, but does not show strong evidence of control by risk-mediating DHS clusters (*γ* = 0.12). The protective MS effect in this region is associated with an increase in *CD58* expression, leading to an up-regulation of the transcription factor *FoxP3* via *CD2.* This results in enhanced functioning of CD4+CD25^high^ regulatory T cells, thought to be defective in MS patients ^26^. Similarly, we are able to prioritize *ETS2* for an inflammatory bowel disease association (IBD) on chromosome 21 (*γ* = 0.92, Supplementary Figure 6), *CD40* for another MS association on chromosome 20 (*γ* = 0.31, Supplementary Figure 7), and *IRF8* for a rheumatoid arthritis (RA) association on chromosome 16 (*γ* = 0.43, Supplementary Figure 8).

**Figure 2:**
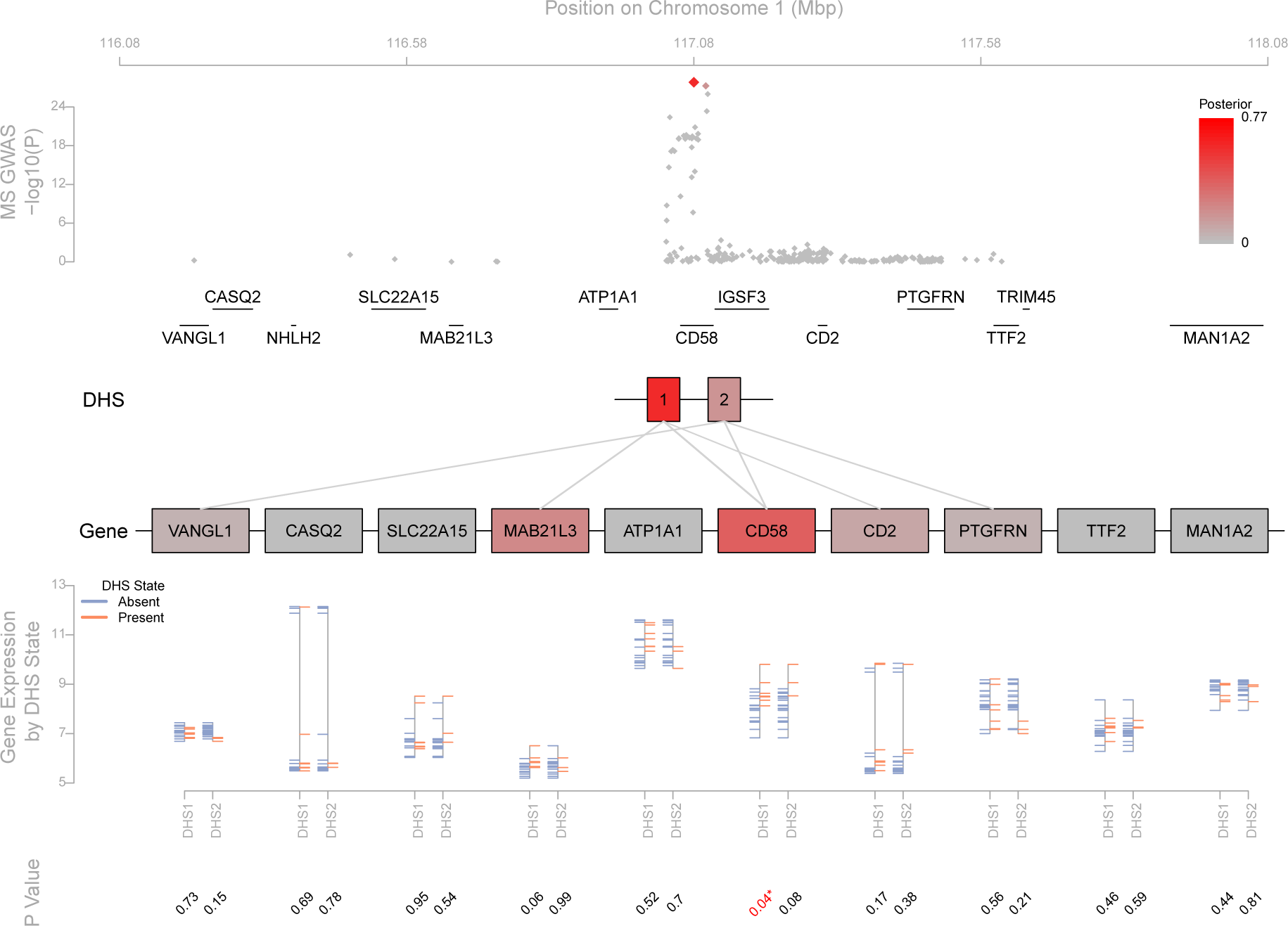
Two DHS clusters identify changes to CD58 regulation as mediating multiple sclerosis risk on chromosome 1. (A) A strong association peak on chromosome 1 localizes to the *CD58* locus. (B) Credible interval mapping identifies two DHS clusters explaining a total of 98.8% of the posterior probability of association. (C) The accessibility of these DHS clusters is correlated to expression levels of several genes in the region. By partitioning the posterior probability of association attributable to each DHS cluster by the strength of this correlation, we find that 48.2% can be attributed to *CD58,* with the next-highest scoring genes *MAB21L3* and *CD2* being attributed 27.2% and 12% respectively. (D) Note that the correlation between DHS1 accessibility and *CD58* expression level is particularly strong across tissues.

Many Immunochip loci harbor associations to multiple diseases, suggesting that a portion of risk is shared^27,28^. Consistent with this observation, we found that 24 Immunochip loci had ρ > 0.25 for more than one disease, representing 59 of the 301 initially considered associations. Of these, 17/24 loci showed regulatory potential in two AID, with four, two, and a single locus showing regulatory potential to three, four and five AID, respectively. Due to the correlation imposed by linkage disequilibrium, it remains challenging to conclude that associations to different traits in the same locus represent a true shared effect, where the same underlying causal variant drives risk for multiple diseases^29^. We therefore sought to establish if associations to different diseases in these 24 loci identify the same DHS clusters and prioritize the same genes, indicating a shared effect. We found striking examples of shared and distinct effects across these 24 loci. For example, five diseases show association to a region of chromosome 6, with the most significant SNPs residing in the coding region of *BACH2.* We are able to prioritize the associations for autoimmune thyroid disease (AITD), MS and type I diabetes (T1D) to a single DHS cluster each, and independently prioritize *MDN1* as the most likely target gene for these effects *(γ_AIT_D* = 0.81, *γ_MS_* = 0.42, and *γ_T_*_1_ *D* = 0.73, Figure 3). Our model only attributes a small proportion of the overall posterior probability of association to *BACH2* for these diseases (*γ_AIT_ D* = 0.14, *γ*_MS_ = 0.09, and *γ_T_*_1_*D* = 0.13). In contrast, we find that the associations for IBD and celiac disease (CEL) each identify different DHS clusters and prioritize *MAP3K7* (*γ_AIT_ D* = 0.59) and *GABBR2* (*γ_CEL_* = 0.13), respectively, despite the credible intervals for these diseases essentially overlapping those for AITD, MS and T1D (Figure 3). We note that the most associated SNPs for MS, AITD, and T1D are the same (rs72928038), and the *R*^2^ between this SNP and the most associated SNPs of IBD (rs1847472) and CEL (rs7753008) are 0.34 and 0.25, respectively. Similarly, we identify the same DHS cluster and prioritize *CTLA4* (*γ_AIT_ D* = 0.47, *γ_RA_* = 0.38, and *γ_T_*_1_ *D* = 0.52) and *ICOS* (*γ_AIT_ D* = 0.41, *γ_RA_* = 0.33, and *γ_T_*_1_ *D* = 0.46) for AITD, RA and T1D associations on chromosome 2 (Supplementary Figure 9). We are thus able to begin resolving associations across multiple diseases into shared and distinct effects in the same locus.

**Figure 3:**
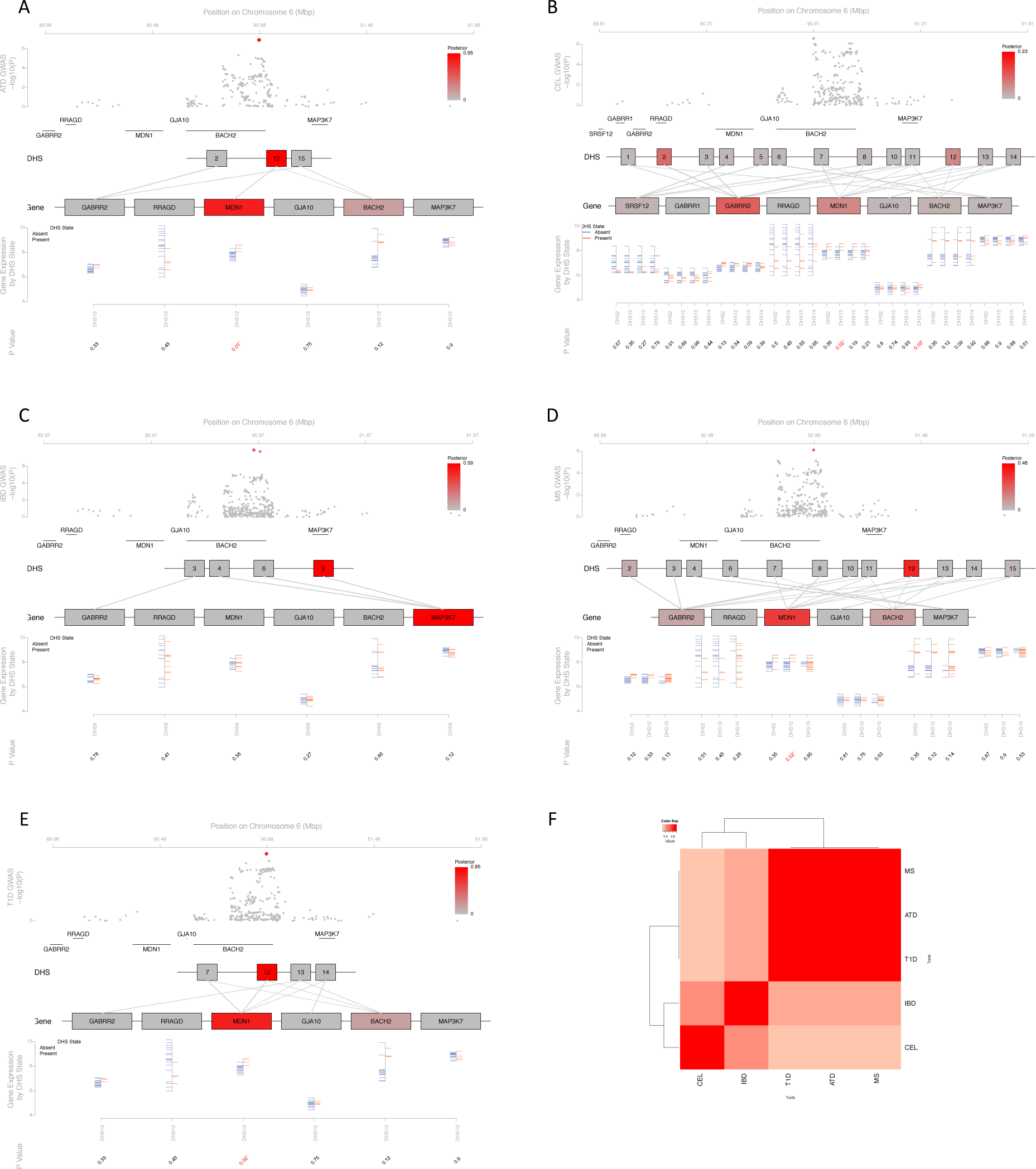
A locus on chromosome 6 harboring association to five diseases identifies MDN1, not BACH2, as driving risk to autoimmune thyroid disease, multiple sclerosis and type 1 diabetes. Association to (A) autoimmune thyroid disease; (B) celiac diease; (C) inflammatory bowel disease; (D) multiple sclerosis; and (E) type 1 diabetes localizes over the coding region of *BACH2.* However, we can attribute the majority of regulatory potential to a single DHS cluster (DHS12) in (A), (D) and (E), which is correlated to the expression of *MDN1,* encoded > 500*kb* from the most associated variant. We find a much weaker correlation to *BACH2* expression. In celiac disease, DHS12 receives the second-highest *ρ* in the locus, but *GABBR2* receives a higher overall posterior *(γ_MDN_*_1_ = 0.074 and *γ_GABBR2_* = 0.134, respectively). In contrast, the strongest IBD posterior is attributed to DHS9, which implicates *MAP3K7.* (F) These differences are consistent with differing LD levels between the most associated SNPs for each disease in the region.

To more generally assess how our approach resolves shared associations, we assessed the overlap between shared signals in the 24 loci. We compared the overlap between 51 pairs of associations in terms of most associated markers, credible interval sets, DHS clusters harboring CI variants, and genes identified (Table 3). We found that, whilst the overlap between lead variants was low, we could more often identify the same DHS clusters and prioritize the same genes (Fisher exact test between proportion of lead SNPs and prioritized genes *p* = 0.014). We found the rate of prioritized gene overlap is correlated to linkage disequilibrium between lead variants (Supplementary Figure 10), suggesting that though GWAS may not identify the same variant representing a shared association, shared effects can clearly be identified by considering the likely functional effects in a locus. These observations hold true when we only consider the 17 loci harboring two disease associations (Supplementary Table 3 and Supplementary Figure 10), indicating our conclusions are not based on biases in a minority of loci harboring many associations. Thus, our approach can uncover biological pleiotropy^30^ across diseases even when the identity of the causal variant remains unknown, beyond the comparison of credible interval sets.

**Table 3:** regulatory fine-mapping resolves associations to multiple diseases in the same loci. When different diseases are associated to the same locus, we find that lead SNPs are often different, credible interval sets and DHS clusters within those sets overlap to a greater extent, and regulatory fine mapping prioritizes the same genes across diseases. Thus, identifying risk-mediating genes partially overcomes the limited resolution of fine-mapping due to linkage disequilibrium.

|  | Concordance | Discordance | Jaccard Coefficient |
| --- | --- | --- | --- |
| **Most Associated SNPs** | 6 | 45 | 0.12 |
| **CI SNPs (mean)** | 6.8 | 31.16 | 0.21 |
| **Prioritized CI SNPs (mean)** | 2.2 | 9.47 | 0.25 |
| **Prioritized DHS Clusters (mean)** | 2.47 | 8.51 | 0.27 |
| **Prioritized Genes** | 16 | 35 | 0.31 |

We have described an approach to detect gene regulatory regions driving disease risk and through them, the genes likely to mediate pathogenesis, through robust re-analysis of public data. We find substantial evidence of regulatory potential in a substantial proportion (44%) of loci across nine AID, and resolve these to a single gene in 104/132 (79%) controlled by regulatory regions active in immune cells. In the majority of loci we examine, we do not prioritize the gene closest to the maximally associated marker. This suggests that risk-mediating regulatory elements act at considerable distances, either by influencing the overall transcriptional landscape of the region or by acting on individual genes at a distance through DNA looping events mediated by DNA-protein interactions^31^. These competing explanations make different predictions: the former implies many genes will be controlled by the risk-mediating regulator, whereas the latter predicts a limited number of targets. As we are able to prioritize a single gene in the majority of cases, our results strongly suggest that risk is mediated by changes to specific gene regulatory programs affecting particular genes, which must be involved in pathogenesis.

More broadly, the observation that most common, complex disease risk aggregates in gene regulatory regions^3,4,5^ has made the translation of genetic association results into molecular and cellular mechanisms challenging. Fine-mapping is limited in resolution by linkage disequilibrium, making association data alone insufficient to identify a causal variant driving risk in a locus. For example, in a recent Immunochip study of multiple sclerosis^32^, we were able to reduce 14/66 (21%) Immunochip regions to 90% credible interval sets of fewer than 15 variants, and 5/66 to fewer than 5 variants, though increases in sample size will raise the resolution of these approaches^14^. Unlike coding variants, inferring function of non-coding polymorphisms remains challenging, though efforts to integrate functional genomics and population genetics data into composite functional scores^33,34^ or integrating genetic and epigenetic data^11^ are gaining some traction on this problem. Our own work complements these efforts by focusing on identifying individual regulators and the genes they control to generate testable hypotheses of the molecular basis of disease mechanism.

## Methods

### DNase I Hypersensitivity data peak-calling, clustering, and quality control

We obtained processed DNase I hypersensitivity (BED format) sequencing reads for 350 Roadmap Epigenomics Project (REP) samples^15,16^ corresponding to 73 cell types from http://www.genboree.org/EdaccData/Current-Release/experiment-sample/Chromatin_Accessibility/. For each sample, we called 150bp DNase I hypersensitive sites (DHS) passing a 1% FDR threshold^17^. We found 56 tissues with at least two replicates, which our statistical replication design requires, and limited our analysis to these. Where more than two replicates were available, we chose the two replicates with the smallest Jaccard distance between their DHS peaks positions on the genome.

To identify corresponding DHS across samples, we calculated the overlap between neighboring peaks across the 112 replicate samples as:

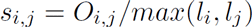

where, *O_i_*,_*j*_ is the number of base pairs shared by DHS *i* and *j*, and *l_i_* and *l_j_* are the length of DHS *i* and *j* respectively. We then grouped DHS with a graph-based approach, the Markov Clustering Algorithm^35^ (MCL) using the default parameters, and defined the coordinates of a DHS cluster as the extreme positions covered by DHS peaks included in that cluster. Finally, we define each cluster as accessible in a sample if we observe at least one DHS peak within its boundaries in that sample.

Both peak calling and MCL clustering are naive to sample labels, so we can test for evidence that DHS clusters replicate in this analysis. We expect that DHS clusters representing true regulatory regions should be consistently accessible or unaccessible in replicate samples. We can thus calculate a replication statistic for DHS cluster *d* as:

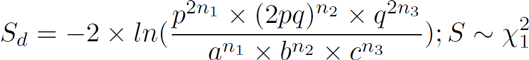

where *n*_1_ is the number of cell types where DHS cluster *d* is active in both replicates; *n*_2_ is the number of cell types where the cluster is active in only one of the two replicates; and *n*_3_ is the number of cell types where the cluster is inactive in both replicates. For *N* = 56 tissues in our data *a* = *n*_1_/*N*, *b* = *n*_2_/*N* and *c* = *n*_3_/*N*. Further, if r is the number of samples where DHS cluster is active, then *p* = *r*/(2 × *N*), and *q* is 1 — *p*. Note that we distinguish between the number of cell types (*N* = 56) and number of samples considered (2 × *N* = 112). We expect S_d_ to follow a distribution, and selected DHS clusters passing a nominal significance threshold of *p_d_* ≤ 0.05. To assess if DHS clusters capture the majority of disease-relevant signal, we compared the proportion of disease heritability (*h*^2^*g*) explained by all DHS detected peaks in a tissue to that explained by the active DHS clusters we annotated^6^. For this we used genome-wide association summary statistics for MS ^36^ and IBD^37^.

### Expression profile processing and analysis across REP tissues

There are 88 Roadmap Epigenomics Project (REP) samples corresponding to 27 cell types profiled on the Affymetrix HuEx-1_0-st-v2 exon array, which we downloaded as raw CEL files on 9/25/2013 from http://www.genboree.org/EdaccData/Current-Release/experiment-sample/Expression_Array/. We processed these data using standard methods available from the BioConductor project^38^. Briefly, we filtered cross-hybridizing probesets, corrected background intensities with RMA and quantile normalizated the remaining probeset intensities across samples. We then collapsed probesets to transcript-level intensities, and mapped transcripts to genes using the current Gencode annotations for human genes (version 12), removing any transcripts without a single exact match to a gene annotation. We then identified the 22 tissues with matched DHS data, averaged measurements over all replicates of each tissue, and quantile normalized the resulting dataset, comprising 13822 transcripts mapping to 13771 unique geneIDs.

### Credible interval mapping for Immunochip loci

We obtained publicly available summary association statistics from case/control cohorts profiled on the Immunochip (Immunobase, http://www.immunobase.org; accessed May 2015) for autoimmune thyroid disease (ATD)^39^, celiac disease (CEL)^40^, inflammatory bowel disease (IBD)^41^, juvenile idiopathic arthritis (JIA)^42^, multiple sclerosis (MS)^32^, primary biliary cirrhosis (PBC)^43^, psoriasis (PSO)^44^, rheumatoid arthritis (RA)^45^, and type 1 diabetes (T1D)^46^ (Table 1). For each of the nine diseases, we compiled a list of genome-wide significant associations from the largest published GWAS^39,40,32,47,46,42,44,37,45^. We then pruned this list of lead SNPs to include only those that overlap densely genotyped regions of Immunochip data and were present in the 1000 Genomes European ancestry cohorts^48^. We excluded the Major Histocompatibility Complex (MHC) region on chromosome 6, where fine-mapping has been previously reported^20^. As summary statistics for conditional associations are not available, we limited our analyses to primary reported signals in each disease.

We identified credible interval SNPs explaining 99% of the posterior probability of association for the remaining lead SNPs^13,14^. For each lead SNP, we identified SNPs within 2Mb in linkage disequilibrium *r*^2^ ≥ 0.1 in the non-Finnish European 1000 Genomes reference panels^48^. For each set *S* of these SNPs, we calculated posterior probabilities of association as

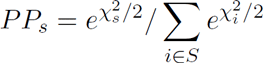

where 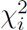 is the Immunochip association chi-square test statistics of SNP *i*. We then selected the smallest number of SNPs required to explain 99% of the posterior probability.

### Calculating regulatory potential of disease loci

We first overlapped credible interval (CI) SNPs with our DHS clusters, then computed the posterior probability of association attributable to each DHS cluster *d* as

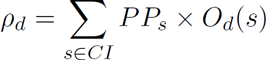

where *PP_s_* is the posterior probability of association for SNP *s*. *O_d_*(*s*) is equal to one if SNP *s* is located on DHS cluster *d* or the 100 bp flanking region each side of DHS cluster *d*, and it is zero otherwise. For SNPs overlapping two or more DHS clusters or their 100 bp flanking regions, we divided its posterior probability *PP_s_* between those DHS clusters equally. We then calculated the regulatory potential of each disease risk locus over all DHSs in the locus as

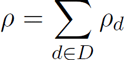

where *D* is the set of all DHS clusters in the region.

### Calculating posterior probabilities of association for each gene in a risk locus

We identified all genes within 1Mb of the lead SNP for each locus, and for all DHS clusters with (*ρ_d_* > 0), computed the correlation between transcript levels and DHS accessibility across the 22 REP tissues with a two-sided Wilcoxon rank sum test. To account for the correlation structure between expression levels, we estimated the expected null empirically. We first decorrelate the matrix of gene expression levels to (*W_P_CA*) using PCA whitening, then use the Cholesky decomposition of the covariance matrix (*L*) to obtain the expected null as *G_Null_* = *L'W_P_CA* (Supplementary Figures 11 and 12). For any given DHS cluster *d*, we computed the Wilcoxon rank sum test statistics between d and all genes of *G_Null_*. This formed our null Wilcoxon rank sum test statistics (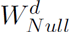). From this null, we computed empirical P-values of significance of correlation between DHS cluster *d* and gene *g* as

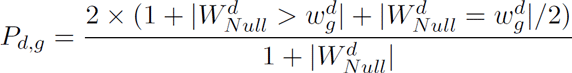

Where 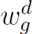 is the Wilcoxon rank sum test statistics between DHS cluster *d* and gene *g*, and |·| denotes the number of events satisfying the enclosed criterion. This formulation accounts for the two-sided test. We used a permutation-based approach to assess the significance of the correlation between DHS clusters and gene expression using a random set of 2000 genes from across the genome. We correlated each random gene to each DHS cluster, and compared test genes against this expected distribution of correlation coefficients to obtain an empirical P value (Supplementary Figure 12).

We next calculated the proportion of posterior probability of association transmitted from DHS cluster *d* to gene *g* as

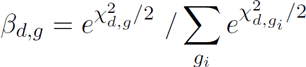

where 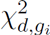 is the chi-squared test statistic corresponding to the empirical correlation *P* value for DHS cluster *d* and gene *g_i_*. From this we computed the total posterior transmitted from DHS cluster *d* to gene *g* as

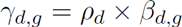

For each gene, we then sum over all DHS clusters *D* to obtain the overall posterior probability of association:

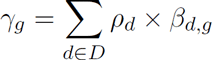

In practice, if *P_d_*,_*g*_ > 0.25 we set *β_d_*,_*g*_ to zero to control noise from small values (Supplementary Figure 13)

### Enrichment of allele—specific accessibility, tissue specificity and functional class for DHS clusters

From Maurano *et. al*.^22^ we obtained a list of 64,597/362,284 SNPs across the genome associated to allele-specific DHS accessibility in heterozygous individuals at 5% FDR. For each disease, we calculated if credible interval SNPs overlapping DHS are likelier to show allelic imbalance than expected by chance using Fisher’s exact test. We also calculated this for all CI SNPs from all diseases as a joint set. We found this enrichment to be consistent across minor allele frequency bins (Supplementary Figure 14).

We used Fisher’s exact test to determine if DHS clusters harboring credible interval SNPs are preferentially active in each tissue. For each tissue, we compare the proportion of active DHS clusters to the genome-wide expectation of active DHS clusters in that tissue. We used the same process to determine enrichment for functional categories defined by ChromHMM ?, and identified genomic functions of DHS clusters through overlapping them with annotated ChromHMM regions (Supplementary Figure 5).

## Acknowledgements

C. C. and P.S. were partly supported by a shared research agreement with Biogen, Inc, who had no role in designing or interpreting this study. We are grateful to the International Multiple Sclerosis Genetics Consortium and the International Inflammatory Bowel Disease Genetics Consortium, and specifically to Mark Daly and Stephan Ripke, for access to GWAS summary statistics from their respective meta-analyses.

## References

1 A. Zhernakova, C. C. van Diemen, and C. Wijmenga. Detecting shared pathogenesis from the shared genetics of immune-related diseases. Nat. Rev. Genet., 10(1):43–55, Jan 2009.

2 L. A. Zenewicz, C. Abraham, R. A. Flavell, and J. H. Cho. Unraveling the genetics of autoimmunity. Cell, 140(6):791–797, Mar 2010.

3 M. T. Maurano, R. Humbert, E. Rynes, R. E. Thurman, E. Haugen, H. Wang, A. P. Reynolds, R. Sandstrom, H. Qu, J. Brody, A. Shafer, F. Neri, K. Lee, T. Kutyavin, S. Stehling-Sun, A. K. Johnson, T. K. Canfield, E. Giste, M. Diegel, D. Bates, R. S. Hansen, S. Neph, P. J. Sabo, S. Heimfeld, A. Raubitschek, S. Ziegler, C. Cotsapas, N. Sotoodehnia, I. Glass, S. R. Sunyaev, R. Kaul, and J. A. Stamatoyannopoulos. Systematic localization of common disease-associated variation in regulatory DNA. Science, 337(6099):1190–1195, Sep 2012.

4 K. K. Farh, A. Marson, J. Zhu, M. Kleinewietfeld, W. J. Housley, S. Beik, N. Shoresh, H. Whitton, R. J. Ryan, A. A. Shishkin, M. Hatan, M. J. Carrasco-Alfonso, D. Mayer, C. J. Luckey, N. A. Patsopoulos, P. L. De Jager, V. K. Kuchroo, C. B. Epstein, M. J. Daly, D. A. Hafler, and B. E. Bernstein. Genetic and epigenetic fine mapping of causal autoimmune disease variants. Nature, 518(7539):337–343, Feb 2015.

5 A. Gusev and et. al. Partitioning heritability of regulatory and cell-type-specific variants across 11 common diseases. Am. J. Hum. Genet., 95(5):535–552, Nov 2014.

6 H. K. Finucane, B. Bulik-Sullivan, A. Gusev, G. Trynka, Y. Reshef, P. R. Loh, V. Anttila, H. Xu, C. Zang, K. Farh, S. Ripke, F. R. Day, S. Purcell, E. Stahl, S. Lindstrom, J. R. Perry, Y. Okada, S. Raychaudhuri, M. J. Daly, N. Patterson, B. M. Neale, and A. L. Price. Partitioning heritability by functional annotation using genome-wide association summary statistics. Nat. Genet., 47(11):1228–1235, Nov 2015.

7 G. Trynka, C. Sandor, B. Han, H. Xu, B. E. Stranger, X. S. Liu, and S. Raychaudhuri. Chromatin marks identify critical cell types for fine mapping complex trait variants. Nat. Genet., 45(2):124–130, Feb 2013.

8 K. J. Karczewski, J. T. Dudley, K. R. Kukurba, R. Chen, A. J. Butte, S. B. Montgomery, and M. Snyder. Systematic functional regulatory assessment of disease-associated variants. Proc. Natl. Acad. Sci. U.S.A., 110(23):9607–9612, Jun 2013.

9 O. Corradin, A. Saiakhova, B. Akhtar-Zaidi, L. Myeroff, J. Willis, R. Cowper-Sal lari, M. Lupien, S. Markowitz, and P. C. Scacheri . Combinatorial effects of multiple enhancer variants in linkage disequilibrium dictate levels of gene expression to confer susceptibility to common traits. Genome Res., 24(1):1–13, Jan 2014.

10 J. Z. Liu, M. A. Almarri, D. J. Gaffney, G. F. Mells, L. Jostins, H. J. Cordell, S. J. Ducker, D. B. Day, M. A. Heneghan, J. M. Neuberger, P. T. Donaldson, A. J. Bathgate, A. Burroughs, M. H. Davies, D. E. Jones, G. J. Alexander, J. C. Barrett, R. N. Sandford, C. A. Anderson, G. Alexander, A. Bathgate, A. Burroughs, H. Cordell, M. Davies, P. Donaldson, M. Heneghan, D. Jones, G. Mells, J. Neuberger, C. Thain, R. Sandford, B. Street, C. Lye, C. Lai, T. Yapp, R. Sturgess, C. Healey, M. Czajkowski, S. Peter, J. Thornton, S. Mann, K. Kapur, R. Marley, G. Foster, J. Ramage, R. Harvey, N. MacDougall, C. J. Shorrock, G. Lipscomb, P. Southern, N. Parnell, J. Tibble, D. Gorard, G. Mells, M. Dawwas, R. Aspinall, S. Dolwani, M. Foxton, H. Mitchison, I. Gooding, M. Patel, R. Ede, A. Austin, R. Dawood, J. Sayer, C. Hovell, N. Fisher, M. Carter, K. Koss, A. Piotrowicz, D. Banait, D. Neal, G. Lim, A. Ala, A. Saeed, J. Brown, S. Thomas, M. Wilkinson, J. Ridpath, T. Ngatchu, S. Levi, R. Ransford, R. Dickinson, R. Shidrawi, G. Abouda, I. Rees, I. Salam, F. Ali, M. Narain, A. Brown, S. Khakoo, S. Williams, M. Williams, A. Chilton, R. Westbrook, M. Heneghan, C. Rodrigues, M. Davies, M. Aldersley, C. Millson, S. Sen, G. Bird, L. Smith, K. Yoong, N. Rajendran, R. Mathew, G. MacFaul, A. Shah, C. Evans, S. Saha, P. Bramley, A. Fraser, P. Mills, T. Shallcross, D. de Las Heras, C. Sheen, R. Crofton, A. Prach, A. Shepherd, H. Kennedy, S. Rushbrook, R. Przemioslo, C. McDonald, B. Javaid, B. Chaudhury, J. Metcalf, D. Ramanaden, J. Gasem, R. Evans, U. Shmueli, A. Naqvi, J. Collier, H. Klass, M. Ninkovic, M. Cramp, P. Goggin, B. Hoeroldt, G. Lipscomb, E. Williams, H. Hussaini, R. Devon, R. Ayres, J. Makanyanga, A. Burroughs, P. Richardson, M. Lombard, D. Robertson, M. Farrant, A. Tanner, S. Singhal, S. Babu, D. Gleeson, J. Butterworth, K. George, H. Curtis, A. McNair, I. Nasr, A. Douglas, J. Shearman, K. Nash, M. Wright, G. Bray, J. Mclindon, D. Das, G. Whatley, S. Lean, N. Sivaramakrishnan, S. Ducker, D. Jones, D. Preston, A. Douds, M. Brookes, V. S. Wong, S. Pereira, M. Carbone, J. Neuberger, G. Watts, F. Gordon, E. Unitt, A. Grant, M. Cox, S. Whalley, J. Fraser, A. Li, A. Bell, H. Gordon, A. Singhal, I. Ahmad, L. NHS, Y. Ang, J. Gotto, A. Turnbull, C. A. Anderson, J. C. Barrett, J. A. Floyd, C. S, R. McGinnis, N. Soranzo, J. Sambrook, J. Stephens, W. H. Ouwehand, W. L. McArdle, S. M. Ring, D. P. Strachan, G. Alexander, J. C. Barrett, C. M. Bulik, P. J. Conlon, A. Dominiczak, A. Duncanson, A. Hill, G. Lord, A. P. Maxwell, L. Morgan, L. Peltonen, R. N, N. Sheerin, N. Soranzo, F. O. Vannberg, J. C. Barrett, P. Concannon, E. Gray, S. E. Hunt, C. Langford, S. Potter, S. Rich, and D. Simpkin. Dense fine-mapping study identifies new susceptibility loci for primary biliary cirrhosis. Nat. Genet., 44(10):1137–1141, Oct 2012.

11 G. Kichaev and B. Pasaniuc. Leveraging Functional-Annotation Data in Trans-ethnic Fine-Mapping Studies. Am. J. Hum. Genet., 97(2):260–271, Aug 2015.

12 M. A. Schaub, A. P. Boyle, A. Kundaje, S. Batzoglou, and M. Snyder. Linking disease associations with regulatory information in the human genome. Genome Res., 22(9):1748–1759, Sep 2012.

13 J. B. Maller, G. McVean, J. Byrnes, D. Vukcevic, K. Palin, Z. Su, J. M. Howson, A. Auton, S. Myers, A. Morris, M. Pirinen, M. A. Brown, P. R. Burton, M. J. Caulfield, A. Compston, M. Farrall, A. S. Hall, A. T. Hattersley, A. V. Hill, C. G. Mathew, M. Pembrey, J. Satsangi, M. R. Stratton, J. Worthington, N. Craddock, M. Hurles, W. Ouwehand, M. Parkes, N. Rahman, A. Duncanson, J. A. Todd, D. P. Kwiatkowski, N. J. Samani, S. C. Gough, M. I. McCarthy, P. Deloukas, P. Donnelly, J. Aerts, T. Ahmad, H. Arbury, A. Attwood, A. Auton, S. G. Ball, A. J. Balmforth, C. Barnes, J. C. Barrett, I. Barroso, A. Barton, A. J. Bennett, S. Bhaskar, K. Blaszczyk, J. Bowes, O. J. Brand, P. S. Braund, F. Bredin, G. Breen, M. J. Brown, I. N. Bruce, J. Bull, O. S. Burren, J. Burton, J. Byrnes, S. Caesar, N. Cardin, C. M. Clee, A. J. Coffey, J. M. Connell, D. F. Conrad, J. D. Cooper, A. F. Dominiczak, K. Downes, H. E. Drummond, D. Dudakia, A. Dunham, B. Ebbs, D. Eccles, S. Edkins, C. Edwards, A. Elliot, P. Emery, D. M. Evans, G. Evans, S. Eyre, A. Farmer, I. N. Ferrier, E. Flynn, A. Forbes, L. Forty, J. A. Franklyn, T. M. Frayling, R. M. Freathy, E. Giannoulatou, P. Gibbs, P. Gilbert, K. Gordon-Smith, E. Gray, E. Green, C. J. Groves, D. Grozeva, R. Gwilliam, A. Hall, N. Hammond, M. Hardy, P. Harrison, N. Hassanali, H. Hebaishi, S. Hines, A. Hinks, G. A. Hitman, L. Hocking, C. Holmes, E. Howard, P. Howard, J. M. Howson, D. Hughes, S. Hunt, J. D. Isaacs, M. Jain, D. P. Jewell, T. Johnson, J. D. Jolley, I. R. Jones, L. A. Jones, G. Kirov, C. F. Langford, H. Lango-Allen, G. M. Lathrop, J. Lee, K. L. Lee, C. Lees, K. Lewis, C. M. Lindgren, M. Maisuria-Armer, J. Maller, J. Mansfield, J. L. Marchini, P. Martin, D. C. Massey, W. L. McArdle, P. McGuffin, K. E. McLay, G. McVean, A. Mentzer, M. L. Mimmack, A. E. Morgan, A. P. Morris, C. Mowat, P. B. Munroe, S. Myers, W. Newman, E. R. Nimmo, M. C. O'Donovan, A. Onipinla, N. R. Ovington, M. J. Owen, K. Palin, A. Palotie, K. Parnell, R. Pearson, D. Pernet, J. R. Perry, A. Phillips, V. Plagnol, N. J. Prescott, I. Prokopenko, M. A. Quail, S. Rafelt, N. W. Rayner, D. M. Reid, A. Renwick, S. M. Ring, N. Robertson, S. Robson, E. Russell, D. St Clair, J. G. Sambrook, J. D. Sanderson, S. J. Sawcer, H. Schuilenburg, C. E. Scott, R. Scott, S. Seal, S. Shaw-Hawkins, B. M. Shields, M. J. Simmonds, D. J. Smyth, E. Somaskantharajah, K. Spanova, S. Steer, J. Stephens, H. E. Stevens, K. Stirrups, M. A. Stone, D. P. Strachan, Z. Su, D. P. Symmons, J. R. Thompson, W. Thomson, M. D. Tobin, M. E. Travers, B. Turnbull, D. Vukcevic, L. V. Wain, M. Walker, N. M. Walker, C. Wallace, M. Warren-Perry, N. A. Watkins, J. Webster, M. N. Weedon, A. G. Wilson, M. Woodburn, B. P. Wordsworth, C. Yau, A. H. Young, E. Zeggini, M. A. Brown, P. R. Burton, M. J. Caulfield, A. Compston, M. Farrall, S. C. Gough, A. S. Hall, A. T. Hattersley, A. V. Hill, C. G. Mathew, M. Pembrey, J. Satsangi, M. R. Stratton, J. Worthington, M. E. Hurles, A. Duncanson, W. H. Ouwehand, M. Parkes, N. Rahman, J. A. Todd, N. J. Samani, D. P. Kwiatkowski, M. I. McCarthy, N. Craddock, P. Deloukas, and P. Donnelly. Bayesian refinement of association signals for 14 loci in 3 common diseases. Nat. Genet., 44(12):1294–1301, Dec 2012.

14 H. Huang and et. al. Association mapping of inflammatory bowel disease loci to single variant resolution. BioRxiv, 2015.

15 B. E. Bernstein, J. A. Stamatoyannopoulos, J. F. Costello, B. Ren, A. Milosavljevic, A. Meissner, M. Kellis, M. A. Marra, A. L. Beaudet, J. R. Ecker, P. J. Farnham, M. Hirst, E. S. Lander, T. S. Mikkelsen, and J. A. Thomson. The NIH Roadmap Epigenomics Mapping Consortium. Nat. Biotechnol., 28(10):1045–1048, Oct 2010.

16 A. Kundaje, W. Meuleman, J. Ernst, M. Bilenky, A. Yen, A. Heravi-Moussavi, P. Kheradpour, Z. Zhang, J. Wang, M. J. Ziller, V. Amin, J. W. Whitaker, M. D. Schultz, L. D. Ward, A. Sarkar, G. Quon, R. S. Sandstrom, M. L. Eaton, Y. C. Wu, A. R. Pfenning, X. Wang, M. Claussnitzer, Y. Liu, C. Coarfa, R. A. Harris, N. Shoresh, C. B. Epstein, E. Gjoneska, D. Leung, W. Xie, R. D. Hawkins, R. Lister, C. Hong, P. Gascard, A. J. Mungall, R. Moore, E. Chuah, A. Tam, T. K. Canfield, R. S. Hansen, R. Kaul, P. J. Sabo, M. S. Bansal, A. Carles, J. R. Dixon, K. H. Farh, S. Feizi, R. Karlic, A. R. Kim, A. Kulkarni, D. Li, R. Lowdon, G. Elliott, T. R. Mercer, S. J. Neph, V. Onuchic, P. Polak, N. Rajagopal, P. Ray, R. C. Sallari, K. T. Siebenthall, N. A. Sinnott-Armstrong, M. Stevens, R. E. Thurman, J. Wu, B. Zhang, X. Zhou, A. E. Beaudet, L. A. Boyer, P. L. De Jager, P. J. Farnham, S. J. Fisher, D. Haussler, S. J. Jones, W. Li, M. A. Marra, M. T. McManus, S. Sunyaev, J. A. Thomson, T. D. Tlsty, L. H. Tsai, W. Wang, R. A. Waterland, M. Q. Zhang, L. H. Chadwick, B. E. Bernstein, J. F. Costello, J. R. Ecker, M. Hirst, A. Meissner, A. Milosavljevic, B. Ren, J. A. Stamatoyannopoulos, T. Wang, M. Kellis, A. Kundaje, W. Meuleman, J. Ernst, M. Bilenky, A. Yen, A. Heravi-Moussavi, P. Kheradpour, Z. Zhang, J. Wang, M. J. Ziller, V. Amin, J. W. Whitaker, M. D. Schultz, L. D. Ward, A. Sarkar, G. Quon, R. S. Sandstrom, M. L. Eaton, Y. C. Wu, A. Pfenning, X. Wang, M. Claussnitzer, Y. Liu, C. Coarfa, R. A. Harris, N. Shoresh, C. B. Epstein, E. Gjoneska, D. Leung, W. Xie, R. D. Hawkins, R. Lister, C. Hong, P. Gascard, A. J. Mungall, R. Moore, E. Chuah, A. Tam, T. K. Canfield, R. S. Hansen, R. Kaul, P. J. Sabo, M. S. Bansal, A. Carles, J. R. Dixon, K. H. Farh, S. Feizi, R. Karlic, A. R. Kim, A. Kulkarni, D. Li, R. Lowdon, G. Elliott, T. R. Mercer, S. J. Neph, V. Onuchic, P. Polak, N. Rajagopal, P. Ray, R. C. Sallari, K. T. Siebenthall, N. A. Sinnott-Armstrong, M. Stevens, R. E. Thurman, J. Wu, B. Zhang, X. Zhou, N. Abdennur, M. Adli, M. Akerman, L. Barrera, J. Antosiewicz-Bourget, T. Ballinger, M. J. Barnes, C. Bates, R. J. Bell, D. A. Bennett, K. Bianco, C. Bock, P. Boyle, J. Brinchmann, P. Caballero-Campo, R. Camahort, M. J. Carrasco-Alfonso, T. Charnecki, H. Chen, Z. Chen, J. B. Cheng, S. Cho, A. Chu, W. Y. Chung, C. Cowan, Q. Athena Deng, V. Deshpande, M. Diegel, B. Ding, T. Durham, L. Echipare, L. Edsall, D. Flowers, O. Genbacev-Krtolica, C. Gifford, S. Gillespie, D. Giste, I. A. Glass, A. Gnirke, M. Gormley, H. Gu, J. Gu, D. A. Hafler, M. J. Hangauer, M. Hariharan, M. Hatan, E. Haugen, Y. He, S. Heimfeld, S. Herlofsen, Z. Hou, R. Humbert, R. Issner, A. R. Jackson, H. Jia, P. Jiang, A. K. Johnson, T. Kadlecek, B. Kamoh, M. Kapidzic, J. Kent, A. Kim, M. Kleinewietfeld, S. Klugman, J. Krishnan, S. Kuan, T. Kutyavin, A. Y. Lee, K. Lee, J. Li, N. Li, Y. Li, K. L. Ligon, S. Lin, Y. Lin, J. Liu, Y. Liu, C. J. Luckey, Y. P. Ma, C. Maire, A. Marson, J. S. Mattick, M. Mayo, M. McMaster, H. Metsky, T. Mikkelsen, D. Miller, M. Miri, E. Mukamel, R. P. Nagarajan, F. Neri, J. Nery, T. Nguyen, H. O'Geen, S. Paithankar, T. Papayannopoulou, M. Pelizzola, P. Plettner, N. E. Propson, S. Raghuraman, B. J. Raney, A. Raubitschek, A. P. Reynolds, H. Richards, K. Riehle, P. Rinaudo, J. F. Robinson, N. B. Rockweiler, E. Rosen, E. Rynes, J. Schein, R. Sears, T. Sejnowski, A. Shafer, L. Shen, R. Shoemaker, M. Sigaroudinia, I. Slukvin, S. Stehling-Sun, R. Stewart, S. L. Subramanian, K. Suknuntha, S. Swanson, S. Tian, H. Tilden, L. Tsai, M. Urich, I. Vaughn, J. Vierstra, S. Vong, U. Wagner, H. Wang, T. Wang, Y. Wang, A. Weiss, H. Whitton, A. Wildberg, H. Witt, K. J. Won, M. Xie, X. Xing, I. Xu, Z. Xuan, Z. Ye, C. A. Yen, P. Yu, X. Zhang, X. Zhang, J. Zhao, Y. Zhou, J. Zhu, Y. Zhu, S. Ziegler, A. E. Beaudet, L. A. Boyer, P. L. De Jager, P. J. Farnham, S. J. Fisher, D. Haussler, S. J. Jones, W. Li, M. A. Marra, M. T. McManus, S. Sunyaev, J. A. Thomson, T. D. Tlsty, L. H. Tsai, W. Wang, R. A. Waterland, M. Q. Zhang, L. H. Chadwick, B. E. Bernstein, J. F. Costello, J. R. Ecker, M. Hirst, A. Meissner, A. Milosavljevic, B. Ren, J. A. Stamatoyannopoulos, T. Wang, M. Kellis, B. E. Bernstein, J. F. Costello, J. R. Ecker, M. Hirst, A. Meissner, A. Milosavljevic, B. Ren, J. A. Stamatoyannopoulos, T. Wang, and M. Kellis. Integrative analysis of 111 reference human epigenomes. Nature, 518(7539): 317–330, Feb 2015.

17 S. John, P. J. Sabo, R. E. Thurman, M. H. Sung, S. C. Biddie, T. A. Johnson, G. L. Hager, and J. A. Stamatoyannopoulos . Chromatin accessibility pre-determines glucocorticoid receptor binding patterns. Nat. Genet., 43(3):264–268, Mar 2011.

18 A. Cortes and M. A. Brown. Promise and pitfalls of the Immunochip. Arthritis Res. Ther., 13 (1):101, 2011.

19 M. Parkes, A. Cortes, D. A. van Heel, and M. A. Brown. Genetic insights into common pathways and complex relationships among immune-mediated diseases. Nat. Rev. Genet., 14(9): 661–673, Sep 2013.

20 L. Moutsianas, L. Jostins, A. H. Beecham, A. T. Dilthey, D. K. Xifara, M. Ban, T. S. Shah, N. A. Patsopoulos, L. Alfredsson, C. A. Anderson, K. E. Attfield, S. E. Baranzini, J. Barrett, T. M. Binder, D. Booth, D. Buck, E. G. Celius, C. Cotsapas, S. D'Alfonso, C. A. Dendrou, P. Donnelly, B. Dubois, B. Fontaine, L. Lar Fugger, A. Goris, P. A. Gourraud, C. Graetz, B. Hemmer, J. Hillert, I. Kockum, S. Leslie, C. M. Lill, F. Martinelli-Boneschi, J. R. Oksenberg, T. Olsson, A. Oturai, J. Saarela, H. B. S?ndergaard, A. Spurkland, B. Taylor, J. Winkelmann, E. Zipp, J. L. Haines, M. A. Pericak-Vance, C. C. Spencer, G. Stewart, D. A. Hafler, A. J. Ivinson, H. F. Harbo, S. L. Hauser, P. L. De Jager, A. Compston, J. L. McCauley, S. Sawcer, and G. McVean. Class II HLA interactions modulate genetic risk for multiple sclerosis. Nat. Genet., 47(10):1107–1113, Oct 2015.

21 G. Trynka, H. J. Westra, K. Slowikowski, X. Hu, H. Xu, B. E. Stranger, R. J. Klein, B. Han, and S. Raychaudhuri. Disentangling the Effects of Colocalizing Genomic Annotations to Functionally Prioritize Non-coding Variants within Complex-Trait Loci. Am. J. Hum. Genet., 97 (1):139–152, Jul 2015.

22 M. T. Maurano, E. Haugen, R. Sandstrom, J. Vierstra, A. Shafer, R. Kaul, and J. A. Stamatoyannopoulos. Large-scale identification of sequence variants influencing human transcription factor occupancy in vivo. Nat. Genet., 47(12):1393–1401, Dec 2015.

23 T. Cremer and C. Cremer. Chromosome territories, nuclear architecture and gene regulation in mammalian cells. Nat. Rev. Genet., 2(4):292–301, Apr 2001.

24 D. U. Gorkin, D. Leung, and B. Ren. The 3D genome in transcriptional regulation and pluripotency. Cell Stem Cell, 14(6):762–775, Jun 2014.

25 Z. Tang, O. J. Luo, X. Li, M. Zheng, J. J. Zhu, P. Szalaj, P. Trzaskoma, A. Magalska, J. Wlodarczyk, B. Ruszczycki, P. Michalski, E. Piecuch, P. Wang, D. Wang, S. Z. Tian, M. Penrad-Mobayed, L. M. Sachs, X. Ruan, C. L. Wei, E. T. Liu, G. M. Wilczynski, D. Plewczynski, F. Li, and Y. Ruan. CTCF-Mediated Human 3D Genome Architecture Reveals Chromatin Topology for Transcription. Cell, 163(7):1611–1627, Dec 2015.

26 P. L. De Jager, C. Baecher-Allan, L. M. Maier, A. T. Arthur, L. Ottoboni, L. Barcellos, J. L. McCauley, S. Sawcer, A. Goris, J. Saarela, R. Yelensky, A. Price, V. Leppa, N. Patterson, P. I. de Bakker, D. Tran, C. Aubin, S. Pobywajlo, E. Rossin, X. Hu, C. W. Ashley, E. Choy, J. D. Rioux, M. A. Pericak-Vance, A. Ivinson, D. R. Booth, G. J. Stewart, A. Palotie, L. Peltonen, B. Dubois, J. L. Haines, H. L. Weiner, A. Compston, S. L. Hauser, M. J. Daly, D. Reich, J. R. Oksenberg, and D. A. Hafler. The role of the CD58 locus in multiple sclerosis. Proc. Natl. Acad. Sci. U.S.A., 106(13):5264–5269, Mar 2009.

27 D. Ellinghaus, L. Jostins, S. L. Spain, A. Cortes, J. Bethune, B. Han, Y. R. Park, S. Raychaudhuri, J. G. Pouget, M. Hubenthal, T. Folseraas, Y. Wang, T. Esko, A. Metspalu, H. J. Westra, L. Franke, T. H. Pers, R. K. Weersma, V. Collij, M. D'Amato, J. Halfvarson, A. B. Jensen, W. Lieb, F. Degenhardt, A. J. Forstner, A. Hofmann, S. Schreiber, U. Mrowietz, B. D. Juran, K. N. Lazaridis, S. Brunak, A. M. Dale, R. C. Trembath, S. Weidinger, M. Weichenthal, E. Ellinghaus, J. T. Elder, J. N. Barker, O. A. Andreassen, D. P. McGovern, T. H. Karlsen, J. C. Barrett, M. Parkes, M. A. Brown, and A. Franke. Analysis of five chronic inflammatory diseases identifies 27 new associations and highlights disease-specific patterns at shared loci. Nat. Genet., 48(5):510–518, May 2016.

28 C. Cotsapas, B. F. Voight, E. Rossin, K. Lage, B. M. Neale, C. Wallace, G. R. Abecasis, J. C. Barrett, T. Behrens, J. Cho, P. L. De Jager, J. T. Elder, R. R. Graham, P. Gregersen, L. Klareskog, K. A. Siminovitch, D. A. van Heel, C. Wijmenga, J. Worthington, J. A. Todd, D. A. Hafler, S. S. Rich, and M. J. Daly. Pervasive sharing of genetic effects in autoimmune disease. PLoS Genet., 7(8):e1002254, Aug 2011.

29 B. K. Bulik-Sullivan, P. R. Loh, H. K. Finucane, S. Ripke, J. Yang, N. Patterson, M. J. Daly, A. L. Price, B. M. Neale, S. Ripke, B. M. Neale, A. Corvin, J. T. Walters, K. H. Farh, P. A. Holmans, P. Lee, B. Bulik-Sullivan, D. A. Collier, H. Huang, T. H. Pers, I. Agartz, E. Agerbo, M. Albus, M. Alexander, F. Amin, S. A. Bacanu, M. Begemann, R. A. Belliveau, J. Bene, S. E. Bergen, E. Bevilacqua, T. B. Bigdeli, D. W. Black, R. Bruggeman, N. G. Buccola, R. L. Buckner, W. Byerley, W. Cahn, G. Cai, M. J. Cairns, D. Campion, R. M. Cantor, V. J. Carr, N. Carrera, S. V. Catts, K. D. Chambert, R. C. Chan, R. Y. Chen, E. Y. Chen, W. Cheng, E. F. Cheung, S. A. Chong, C. Cloninger, D. Cohen, N. Cohen, P. Cormican, N. Craddock, B. Crespo-Facorro, J. J. Crowley, D. Curtis, M. Davidson, K. L. Davis, F. Degenhardt, J. Del Favero, L. E. DeLisi, D. Demontis, D. Dikeos, T. Dinan, S. Djurovic, G. Donohoe, E. Drapeau, J. Duan, F. Dudbridge, N. Durmishi, P. Eichhammer, J. Eriksson, V. Escott-Price, L. Essioux, A. H. Fanous, M. S. Farrell, J. Frank, L. Franke, R. Freedman, N. B. Freimer, M. Friedl, J. I. Friedman, M. Fromer, G. Genovese, L. Georgieva, E. S. Gershon, I. Giegling, P. Giusti-Rodriguez, S. Godard, J. I. Goldstein, V. Golimbet, S. Gopal, J. Gratten, L. de Haan, C. Hammer, M. L. Hamshere, M. Hansen, T. Hansen, V. Haroutunian, A. M. Hartmann, F. A. Henskens, S. Herms, J. N. Hirschhorn, P. Hoffmann, A. Hofman, M. V. Hollegaard, D. M. Hougaard, M. Ikeda, I. Joa, A. Julia, R. S. Kahn, L. Kalaydjieva, S. Karachanak-Yankova, J. Karjalainen, D. Kavanagh, M. C. Keller, B. J. Kelly, J. L. Kennedy, A. Khrunin, Y. Kim, J. Klovins, J. A. Knowles, B. Konte, V. Kucinskas, Z. A. Kucinskiene, H. Kuzelova-Ptackova, A. K. Kahler, C. Laurent, J. L. Keong, S. Lee, S. E. Legge, B. Lerer, M. Li, T. Li, K. Y. Liang, J. Lieberman, S. Limborska, C. M. Loughland, J. Lubinski, J. Lonnqvist, M. Macek, P. K. Magnusson, B. S. Maher, W. Maier, J. Mallet, S. Marsal, M. Mattheisen, M. Mattingsdal, R. W. McCarley, C. McDonald, A. M. McIntosh, S. Meier, C. J. Meijer, B. Melegh, I. Melle, R. I. Mesholam-Gately, A. Metspalu, P. T. Michie, L. Milani, V. Milanova, Y. Mokrab, D. W. Morris, O. Mors, K. C. Murphy, R. M. Murray, I. Myin-Germeys, B. Muller-Myhsok, M. Nelis, I. Nenadic, D. A. Nertney, G. Nestadt, K. K. Nicodemus, L. Nikitina-Zake, L. Nisenbaum, A. Nordin, E. O'Callaghan, C. O'Dushlaine, F. A. O'Neill, S. Y. Oh, A. Olincy, L. Olsen, J. Van Os, C. Pantelis, G. N. Papadimitriou, S. Papiol, E. Parkhomenko, M. T. Pato, T. Paunio, M. Pejovic-Milovancevic, D. O. Perkins, O. Pietilainen, J. Pimm, A. J. Pocklington, J. Powell, A. Price, A. E. Pulver, S. M. Purcell, D. Quested, H. B. Rasmussen, A. Reichenberg, M. A. Reimers, A. L. Richards, J. L. Roffman, P. Roussos, D. M. Ruderfer, V. Salomaa, A. R. Sanders, U. Schall, C. R. Schubert, T. G. Schulze, S. G. Schwab, E. M. Scolnick, R. J. Scott, L. J. Seidman, J. Shi, E. Sigurdsson, T. Silagadze, J. M. Silverman, K. Sim, P. Slominsky, J. W. Smoller, H. C. So, C. C. Spencer, E. A. Stahl, H. Stefansson, S. Steinberg, E. Stogmann, R. E. Straub, E. Strengman, J. Strohmaier, T. Stroup, M. Subramaniam, J. Suvisaari, D. M. Svrakic, J. P. Szatkiewicz, E. Soderman, S. Thirumalai, D. Toncheva, P. A. Tooney, S. Tosato, J. Veijola, J. Waddington, D. Walsh, D. Wang, Q. Wang, B. T. Webb, M. Weiser, D. D. Wildenauer, N. M. Williams, S. Williams, S. H. Witt, A. R. Wolen, E. H. Wong, B. K. Wormley, J. Q. Wu, H. S. Xi, C. C. Zai, X. Zheng, F. Zimprich, N. R. Wray, K. Stefansson, P. M. Visscher, R. Adolfsson, O. A. Andreassen, D. H. Blackwood, E. Bramon, J. D. Buxbaum, A. D. B?rglum, S. Cichon, A. Darvasi, E. Domenici, H. Ehrenreich, T. Esko, P. V. Gejman, M. Gill, H. Gurling, C. M. Hultman, N. Iwata, A. V. Jablensky, E. G. Jonsson, K. S. Kendler, G. Kirov, J. Knight, T. Lencz, D. F. Levinson, Q. S. Li, J. Liu, A. K. Malhotra, S. A. McCarroll, A. McQuillin, J. L. Moran, P. B. Mortensen, B. J. Mowry, M. M. Nothen, R. A. Ophoff, M. J. Owen, A. Palotie, C. N. Pato, T. L. Petryshen, D. Posthuma, M. Rietschel, B. P. Riley, D. Rujescu, P. C. Sham, P. Sklar, D. St Clair, D. R. Weinberger, J. R. Wendland, T. Werge, M. J. Daly, P. F. Sullivan, and M. C. O'Donovan. LD Score regression distinguishes confounding from polygenicity in genome-wide association studies. Nat. Genet., 47(3):291–295, Mar 2015.

30 N. Solovieff, C. Cotsapas, P. H. Lee, S. M. Purcell, and J. W. Smoller. Pleiotropy in complex traits: challenges and strategies. Nat. Rev. Genet., 14(7):483–495, Jul 2013.

31 L. J. Davison, C. Wallace, J. D. Cooper, N. F. Cope, N. K. Wilson, D. J. Smyth, J. M. Howson, N. Saleh, A. Al-Jeffery, K. L. Angus, H. E. Stevens, S. Nutland, S. Duley, R. M. Coulson, N. M. Walker, O. S. Burren, C. M. Rice, F. Cambien, T. Zeller, T. Munzel, K. Lackner, S. Blakenberg, P. Fraser, B. Gottgens, J. A. Todd, T. Attwood, S. Belz, P. Braund, F. Cambien, J. Cooper, A. Crisp-Hihn, P. Diemert, P. Deloukas, N. Foad, J. Erdmann, A. H. Goodall, J. Gracey, E. Gray, R. G. Williams, S. Heimerl, C. Hengstenberg, J. Jolley, U. Krishnan, H. Lloyd-Jones, I. Lugauer, P. Lundmark, S. Maouche, J. S. Moore, D. Muir, E. Murray, C. P. Nelson, J. Neudert, D. Niblett, K. O'Leary, W. H. Ouwehand, H. Pollard, A. Rankin, C. M. Rice, G. Sager, N. J. Samani, J. Sambrook, G. Schmitz, M. Scholz, L. Schroeder, H. Schunkert, A. C. Syvannen, S. Tennstedt, and C. Wallace. Long-range DNA looping and gene expression analyses identify DEXI as an autoimmune disease candidate gene. Hum. Mol. Genet., 21(2): 322–333, Jan 2012.

32 A. H. Beecham, N. A. Patsopoulos, D. K. Xifara, M. F. Davis, A. Kemppinen, C. Cotsapas, T. S. Shah, C. Spencer, D. Booth, A. Goris, A. Oturai, J. Saarela, B. Fontaine, B. Hemmer, C. Martin, F. Zipp, S. D'Alfonso, F. Martinelli-Boneschi, B. Taylor, H. F. Harbo, I. Kockum, J. Hillert, T. Olsson, M. Ban, J. R. Oksenberg, R. Hintzen, L. F. Barcellos, C. Agliardi, L. Alfredsson, M. Alizadeh, C. Anderson, R. Andrews, H. B. S?ndergaard, A. Baker, G. Band, S. E. Baranzini, N. Barizzone, J. Barrett, C. Bellenguez, L. Bergamaschi, L. Bernardinelli, A. Berthele, V. Biberacher, T. M. Binder, H. Blackburn, I. L. Bomfim, P. Brambilla, S. Broadley, B. Brochet, L. Brundin, D. Buck, H. Butzkueven, S. J. Caillier, W. Camu, W. Carpentier, P. Cavalla, E. G. Celius, I. Coman, G. Comi, L. Corrado, L. Cosemans, I. Cournu-Rebeix, B. A. Cree, D. Cusi, V. Damotte, G. Defer, S. R. Delgado, P. Deloukas, A. di Sapio, A. T. Dilthey, P. Donnelly, B. Dubois, M. Duddy, S. Edkins, I. Elovaara, F. Esposito, N. Evangelou, B. Fiddes, J. Field, A. Franke, C. Freeman, I. Y. Frohlich, D. Galimberti, C. Gieger, P. A. Gourraud, C. Graetz, A. Graham, V. Grummel, C. Guaschino, A. Hadjixenofontos, H. Hakonarson, C. Halfpenny, G. Hall, P. Hall, A. Hamsten, J. Harley, T. Harrower, C. Hawkins, G. Hellenthal, C. Hillier, J. Hobart, M. Hoshi, S. E. Hunt, M. Jagodic, I. Jel?i?, A. Jochim, B. Kendall, A. Kermode, T. Kilpatrick, K. Koivisto, I. Konidari, T. Korn, H. Kronsbein, C. Langford, M. Larsson, M. Lathrop, C. Lebrun-Frenay, J. Lechner-Scott, M. H. Lee, M. A. Leone, V. Leppa, G. Liberatore, B. A. Lie, C. M. Lill, M. Linden, J. Link, F. Luessi, J. Lycke, F. Macciardi, S. Mannisto, C. P. Manrique, R. Martin, V. Martinelli, D. Mason, G. Mazibrada, C. McCabe, I. L. Mero, J. Mescheriakova, L. Moutsianas, K. M. Myhr, G. Nagels, R. Nicholas, P. Nilsson, F. Piehl, M. Pirinen, S. E. Price, H. Quach, M. Reunanen, W. Robberecht, N. P. Robertson, M. Rodegher, D. Rog, M. Salvetti, N. C. Schnetz-Boutaud, F. Sellebjerg, R. C. Selter, C. Schaefer, S. Shaunak, L. Shen, S. Shields, V. Siffrin, M. Slee, P. S. Sorensen, M. Sorosina, M. Sospedra, A. Spurkland, A. Strange, E. Sundqvist, V. Thijs, J. Thorpe, A. Ticca, P. Tienari, C. van Duijn, E. M. Visser, S. Vucic, H. Westerlind, J. S. Wiley, A. Wilkins, J. F. Wilson, J. Winkelmann, J. Zajicek, E. Zindler, J. L. Haines, M. A. Pericak-Vance, A. J. Ivinson, G. Stewart, D. Hafler, S. L. Hauser, A. Compston, G. McVean, P. De Jager, S. J. Sawcer, and J. L. McCauley. Analysis of immune-related loci identifies 48 new susceptibility variants for multiple sclerosis. Nat. Genet., 45(11):1353–1360, Nov 2013.

33 M. Kircher, D. M. Witten, P. Jain, B. J. O'Roak, G. M. Cooper, and J. Shendure. A general framework for estimating the relative pathogenicity of human genetic variants. Nat. Genet., 46(3):310–315, Mar 2014.

34 S. Petrovski, A. B. Gussow, Q. Wang, M. Halvorsen, Y. Han, W. H. Weir, A. S. Allen, and D. B. Goldstein. The Intolerance of Regulatory Sequence to Genetic Variation Predicts Gene Dosage Sensitivity. PLoS Genet., 11(9):e1005492, Sep 2015.

35 A. J. Enright, S. Van Dongen, and C. A. Ouzounis. An efficient algorithm for large-scale detection of protein families. Nucleic Acids Res., 30(7):1575–1584, Apr 2002.

36 N. A. Patsopoulos, F. Esposito, J. Reischl, S. Lehr, D. Bauer, J. Heubach, R. Sandbrink, C. Pohl, G. Edan, L. Kappos, D. Miller, J. Montalban, C. H. Polman, M. S. Freedman, H. P. Hartung, B. G. Arnason, G. Comi, S. Cook, M. Filippi, D. S. Goodin, D. Jeffery, P. O'Connor, G. C. Ebers, D. Langdon, A. T. Reder, A. Traboulsee, F. Zipp, S. Schimrigk, J. Hillert, M. Bahlo, D. R. Booth, S. Broadley, M. A. Brown, B. L. Browning, S. R. Browning, H. Butzkueven, W. M. Carroll, C. Chapman, S. J. Foote, L. Griffiths, A. G. Kermode, T. J. Kilpatrick, J. Lechner-Scott, M. Marriott, D. Mason, P. Moscato, R. N. Heard, M. P. Pender, V. M. Perreau, D. Perera, J. P. Rubio, R. J. Scott, M. Slee, J. Stankovich, G. J. Stewart, B. V. Taylor, N. Tubridy, E. Willoughby, J. Wiley, P. Matthews, F. M. Boneschi, A. Compston, J. Haines, S. L. Hauser, J. McCauley, A. Ivinson, J. R. Oksenberg, M. Pericak-Vance, S. J. Sawcer, P. L. De Jager, D. A. Hafler, and P. I. de Bakker. Genome-wide meta-analysis identifies novel multiple sclerosis susceptibility loci. Ann. Neurol., 70(6):897–912, Dec 2011.

37 J. Z. Liu, S. van Sommeren, H. Huang, S. C. Ng, R. Alberts, A. Takahashi, S. Ripke, J. C. Lee, L. Jostins, T. Shah, S. Abedian, J. H. Cheon, J. Cho, N. E. Daryani, L. Franke, Y. Fuyuno, A. Hart, R. C. Juyal, G. Juyal, W. H. Kim, A. P. Morris, H. Poustchi, W. G. Newman, V. Midha, T. R. Orchard, H. Vahedi, A. Sood, J. J. Sung, R. Malekzadeh, H. J. Westra, K. Yamazaki, S. K. Yang, J. C. Barrett, A. Franke, B. Z. Alizadeh, M. Parkes, T. B K, M. J. Daly, M. Kubo, C. A. Anderson, R. K. Weersma, S. Abedian, C. Abraham, J. P. Achkar, T. Ahmad, R. Alberts, B. Alizadeh, L. Amininejad, A. N. Anathakrishnan, V. Andersen, C. A. Anderson, J. M. Andrews, V. Annese, G. Aumais, L. Baidoo, R. N. Baldassano, T. Balschun, P. A. Bampton, M. Barclay, J. C. Barrett, T. M. Bayless, J. Bethge, C. Bewshea, J. C. Bis, A. Bitton, T. B K, G. Boucher, O. Brain, S. Brand, S. R. Brant, C. Buning, J. H. Cheon, A. Chew, J. H. Cho, I. Cleynen, A. Cohain, R. Cooney, A. Croft, M. J. Daly, M. D'Amato, S. Danese, N. E. Daryani, D. De Jong, K. M. de Lange, M. De Vos, G. Denapiene, L. A. Denson, K. L. Devaney, O. Dewit, R. D'Inca, H. E. Drummond, M. Dubinsky, R. H. Duerr, C. Edwards, D. Ellinghaus, M. Esaki, J. Essers, L. R. Ferguson, E. A. Festen, P. Fleshner, T. Florin, D. Franchimont, A. Franke, K. Fransen, Y. Fuyano, R. Gearry, M. Georges, C. Gieger, J. Glas, P. Goyette, T. Green, A. M. Griffiths, S. L. Guthery, H. Hakonarson, J. Halfvarson, K. Hanigan, T. Haritunians, A. Hart, C. Hawkey, N. K. Hayward, M. Hedl, P. Henderson, G. L. Hold, X. Hu, H. Huang, K. Y. Hui, M. Imielinski, A. Ippoliti, O. Jazayeri, L. Jonaitis, L. Jostins, G. Juyal, R. C. Juyal, R. Kalla, T. H. Karlsen, T. Kawaguchi, N. A. Kennedy, M. A. Khan, W. H. Kim, T. Kitazono, G. Kiudelis, M. Kubo, S. Kugathasan, L. Kupcinskas, C. A. Lamb, A. Latiano, D. Laukens, I. C. Lawrance, J. C. Lee, C. W. Lees, M. Leja, N. Lewis, J. Van Limbergen, P. Lionetti, J. Z. Liu, E. Louis, Y. Luo, G. Mahy, M. M. Malekzadeh, R. Malekzadeh, J. Mansfield, S. Marriott, D. Massey, C. G. Mathew, T. Matsui, D. P. McGovern, V. Midha, R. Milgrom, S. Mirzaei, M. Mitrovic, G. W. Montgomery, S. Motoya, C. Mowat, W. G. Newman, A. Ng, S. C. Ng, S. M. Ng, S. Nikolaus, E. R. Nimmo, K. Ning, M. Nothen, I. Oikonomou, T. R. Orchard,O. Palmieri, M. Parkes, A. Phillips, C. Y. Ponsioen, U. Potocnik, H. Poustchi, N. J. Prescott, D. D. Proctor, G. Radford-Smith, J. F. Rahier, S. Raychaudhuri, M. Regueiro, F. Rieder, J. D. Rioux, S. Ripke, R. Roberts, R. K. Russell, J. D. Sanderson, M. Sans, J. Satsangi, E. E. Schadt, S. Schreiber, L. P. Schumm, R. Scott, M. Seielstad, T. Shah, Y. Sharma, M. S. Silverberg, A. Simmons, L. A. Simms, A. Singh, J. Skieceviciene, A. Sood, S. L. Spain, A. H. Steinhart, J. M. Stempak, L. Stronati, J. J. Sung, Y. Suzuki, J. Sventoraityte, A. Takahashi, M. Takazoe, H. Tanaka, K. M. Taylor, A. ter Velde, E. Theatre, L. Torkvist, M. Tremelling, H. H. Uhlig, H. Vahedi, A. van der Meulen, S. van Sommeren, E. Vasiliauskas, N. T. Ventham, S. Vermeire, H. W. Verspaget, T. Walters, K. Wang, M. H. Wang, R. K. Weersma, Z. Wei, D. Whiteman, C. Wijmenga, D. C. Wilson, J. Winkelmann, R. J. Xavier, T. Yamada, K. Yamazaki, S. Zeissig, B. Zhang, C. K. Zhang, H. Zhang, W. Zhang, H. Zhao, and Z. Z. Zhao. Association analyses identify 38 susceptibility loci for inflammatory bowel disease and highlight shared genetic risk across populations. Nat. Genet., 47(9):979–986, Sep 2015.

38 W. Huber, V. J. Carey, R. Gentleman, S. Anders, M. Carlson, B. S. Carvalho, H. C. Bravo, S. Davis, L. Gatto, T. Girke, R. Gottardo, F. Hahne, K. D. Hansen, R. A. Irizarry, M. Lawrence, M. I. Love, J. MacDonald, V. Obenchain, A. K. Ole?, H. Pages, A. Reyes, P. Shannon, G. K. Smyth, D. Tenenbaum, L. Waldron, and M. Morgan. Orchestrating high-throughput genomic analysis with Bioconductor. Nat. Methods, 12(2):115–121, Feb 2015.

39 J. D. Cooper, M. J. Simmonds, N. M. Walker, O. Burren, O. J. Brand, H. Guo, C. Wallace, H. Stevens, G. Coleman, J. A. Franklyn, J. A. Todd, S. C. Gough, J. Aerts, T. Ahmad, H. Arbury, A. Attwood, A. Auton, S. G. Ball, A. J. Balmforth, C. Barnes, J. C. Barrett,I. Barroso, A. Barton, A. J. Bennett, S. Bhaskar, K. Blaszczyk, J. Bowes, O. J. Brand, P. S. Braund, F. Bredin, G. Breen, M. J. Brown, I. N. Bruce, J. Bull, O. S. Burren, J. Burton, J. Byrnes, S. Caesar, N. Cardin, C. M. Clee, A. J. Coffey, J. M. Connell, D. F. Conrad, J. D. Cooper, A. F. Dominiczak, K. Downes, H. E. Drummond, D. Dudakia, A. Dunham, B. Ebbs, D. Eccles, S. Edkins, C. Edwards, A. Elliot, P. Emery, D. M. Evans, G. Evans, S. Eyre, A. Farmer, I. N. Ferrier, E. Flynn, A. Forbes, L. Forty, J. A. Franklyn, T. M. Frayling, R. M. Freathy, E. Giannoulatou, P. Gibbs, P. Gilbert, K. Gordon-Smith, E. Gray, E. Green, C. J. Groves, D. Grozeva, R. Gwilliam, A. Hall, N. Hammond, M. Hardy, P. Harrison, N. Hassanali, H. Hebaishi, S. Hines, A. Hinks, G. A. Hitman, L. Hocking, C. Holmes, E. Howard, P. Howard, J. M. Howson, D. Hughes, S. Hunt, J. D. Isaacs, M. Jain, D. P. Jewell, T. Johnson, J. D. Jolley, I. R. Jones, L. A. Jones, G. Kirov, C. F. Langford, H. Lango-Allen, G. M. Lathrop, J. Lee, K. L. Lee, C. Lees, K. Lewis, C. M. Lindgren, M. Maisuria-Armer, J. Maller, J. Mansfield, J. L. Marchini, P. Martin, D. C. Massey, W. L. McArdle, P. McGuffin, K. E. McLay, G. McVean, A. Mentzer, M. L. Mimmack, A. E. Morgan, A. P. Morris, C. Mowat, P. B. Munroe, S. Myers, W. Newman, E. R. Nimmo, M. C. O'Donovan, A. Onipinla, N. R. Ovington, M. J. Owen, K. Palin, A. Palotie, K. Parnell, R. Pearson, D. Pernet, J. R. Perry, A. Phillips, V. Plagnol, N. J. Prescott, I. Prokopenko, M. A. Quail, S. Rafelt, N. W. Rayner, D. M. Reid, A. Renwick, S. M. Ring, N. Robertson, S. Robson, E. Russell, D. St Clair, J. G. Sambrook, J. D. Sanderson, S. J. Sawcer, H. Schuilenburg, C. E. Scott, R. Scott, S. Seal, S. Shaw-Hawkins, B. M. Shields, M. J. Simmonds, D. J. Smyth, E. Somaskantharajah, K. Spanova, S. Steer, J. Stephens, H. E. Stevens, K. Stirrups, M. A. Stone, D. P. Strachan, Z. Su, D. P. Symmons, J. R. Thompson, W. Thomson, M. D. Tobin, M. E. Travers, C. Turnbull, D. Vukcevic, L. V. Wain, M. Walker, N. M. Walker, C. Wallace, M. Warren-Perry, N. A. Watkins, J. Webster, M. N. Weedon, A. G. Wilson, M. Woodburn, B. P. Wordsworth, C. Yau, A. H. Young, E. Zeggini, M. A. Brown, P. R. Burton, M. J. Caulfield, A. Compston, M. Farrall, S. C. Gough, A. S. Hall, A. T. Hattersley, A. V. Hill, C. G. Mathew, M. Pembrey, J. Satsangi, M. R. Stratton, J. Worthington, M. E. Hurles, A. Duncanson, W. H. Ouwehand, M. Parkes, N. Rahman, J. A. Todd, N. J. Samani, D. P. Kwiatkowski, M. I. McCarthy, N. Craddock, P. Deloukas, and P. Donnelly. Seven newly identified loci for autoimmune thyroid disease. Hum. Mol. Genet., 21(23):5202–5208, Dec 2012.

40 G. Trynka, K. A. Hunt, N. A. Bockett, J. Romanos, V. Mistry, A. Szperl, S. F. Bakker, M. T. Bardella, L. Bhaw-Rosun, G. Castillejo, E. G. de la Concha, R. C. de Almeida, K. R. Dias, C. C. van Diemen, P. C. Dubois, R. H. Duerr, S. Edkins, L. Franke, K. Fransen, J. Gutierrez, G. A. Heap, B. Hrdlickova, S. Hunt, L. Plaza Izurieta, V. Izzo, L. A. Joosten, C. Langford, M. C. Mazzilli, C. A. Mein, V. Midah, M. Mitrovic, B. Mora, M. Morelli, S. Nutland, C. Nunez, S. Onengut-Gumuscu, K. Pearce, M. Platteel, I. Polanco, S. Potter, C. Ribes-Koninckx, I. Ricano-Ponce, S. S. Rich, A. Rybak, J. L. Santiago, S. Senapati, A. Sood, H. Szajewska, R. Troncone, J. Varade, C. Wallace, V. M. Wolters, A. Zhernakova, B. K. Thelma, B. Cukrowska, E. Urcelay, J. R. Bilbao, M. L. Mearin, D. Barisani, J. C. Barrett, V. Plagnol, P. Deloukas, C. Wijmenga, and D. A. van Heel. Dense genotyping identifies and localizes multiple common and rare variant association signals in celiac disease. Nat. Genet., 43(12): 1193–1201, Dec 2011.

41 L. Jostins, S. Ripke, R. K. Weersma, R. H. Duerr, D. P. McGovern, K. Y. Hui, J. C. Lee, L. P. Schumm, Y. Sharma, C. A. Anderson, J. Essers, M. Mitrovic, K. Ning, I. Cleynen, E. Theatre, S. L. Spain, S. Raychaudhuri, P. Goyette, Z. Wei, C. Abraham, J. P. Achkar, T. Ahmad, L. Amininejad, A. N. Ananthakrishnan, V. Andersen, J. M. Andrews, L. Baidoo, T. Balschun, P. A. Bampton, A. Bitton, G. Boucher, S. Brand, C. Buning, A. Cohain, S. Cichon, M. D'Amato, D. De Jong, K. L. Devaney, M. Dubinsky, C. Edwards, D. Ellinghaus, L. R. Ferguson, D. Franchimont, K. Fransen, R. Gearry, M. Georges, C. Gieger, J. Glas, T. Haritunians, A. Hart, C. Hawkey, M. Hedl, X. Hu, T. H. Karlsen, L. Kupcinskas, S. Kugathasan, A. Latiano, D. Laukens, I. C. Lawrance, C. W. Lees, E. Louis, G. Mahy, J. Mansfield, A. R. Morgan, C. Mowat, W. Newman, O. Palmieri, C. Y. Ponsioen, U. Potocnik, N. J. Prescott, M. Regueiro, J. I. Rotter, R. K. Russell, J. D. Sanderson, M. Sans, J. Satsangi, S. Schreiber, L. A. Simms, J. Sventoraityte, S. R. Targan, K. D. Taylor, M. Tremelling, H. W. Verspaget, M. De Vos, C. Wijmenga, D. C. Wilson, J. Winkelmann, R. J. Xavier, S. Zeissig, B. Zhang, C. K. Zhang, H. Zhao, M. S. Silverberg, V. Annese, H. Hakonarson, S. R. Brant, G. Radford-Smith, C. G. Mathew, J. D. Rioux, E. E. Schadt, M. J. Daly, A. Franke, M. Parkes, S. Vermeire, J. C. Barrett, J. H. Cho, M. Barclay, L. Peyrin-Biroulet, M. Chamaillard, J. F. Colombel, M. Cottone, A. Croft, R. D'Inca, J. Halfvarson, K. Hanigan, P. Henderson, J. P. Hugot, A. Karban, N. A. Kennedy, M. A. Khan, M. Lemann, A. Levine, D. Massey, M. Milla, G. W. Montgomery, S. M. Ng, I. Oikonomou, H. Peeters, D. D. Proctor, J. F. Rahier, R. Roberts, P. Rutgeerts, F. Seibold, L. Stronati, K. M. Taylor, L. Torkvist, K. Ublick, J. Van Limbergen, A. Van Gossum, M. H. Vatn, H. Zhang, W. Zhang, J. M. Andrews, P. A. Bampton, M. Barclay, T. H. Florin, R. Gearry, K. Krishnaprasad, I. C. Lawrance, G. Mahy, G. W. Montgomery, G. Radford-Smith, R. L. Roberts, L. A. Simms, L. Amininijad, I. Cleynen, O. Dewit, D. Franchimont, M. Georges, D. Laukens, H. Peeters, J. F. Rahier, P. Rutgeerts, E. Theatre, A. Van Gossum, S. Vermeire, G. Aumais, L. Baidoo, A. M. Barrie, K. Beck, E. J. Bernard, D. G. Binion, A. Bitton, S. R. Brant, J. H. Cho, A. Cohen, K. Croitoru, M. J. Daly, L. W. Datta, C. Deslandres, R. H. Duerr, D. Dutridge, J. Ferguson, J. Fultz, P. Goyette, G. R. Greenberg, T. Haritunians, G. Jobin, S. Katz, R. G. Lahaie, D. P. McGovern, L. Nelson, S. M. Ng, K. Ning, I. Oikonomou, P. Pare, D. D. Proctor, M. D. Regueiro, J. D. Rioux, E. Ruggiero, L. Schumm, M. Schwartz, R. Scott, Y. Sharma, M. S. Silverberg, D. Spears, A. Steinhart, J. M. Stempak, J. M. Swoger, C. Tsagarelis, W. Zhang, C. Zhang, H. Zhao, J. Aerts, T. Ahmad, H. Arbury, A. Attwood, A. Auton, S. G. Ball, A. J. Balmforth, C. Barnes, J. C. Barrett, I. Barroso, A. Barton, A. J. Bennett, S. Bhaskar, K. Blaszczyk, J. Bowes, O. J. Brand, P. S. Braund, F. Bredin, G. Breen, M. J. Brown, I. N. Bruce, J. Bull, O. S. Burren, J. Burton, J. Byrnes, S. Caesar, N. Cardin, C. M. Clee, A. J. Coffey, J. M. Connell, D. F. Conrad, J. D. Cooper, A. F. Dominiczak, K. Downes, H. E. Drummond, D. Dudakia, A. Dunham, B. Ebbs, D. Eccles, S. Edkins, C. Edwards, A. Elliot, P. Emery, D. M. Evans, G. Evans, S. Eyre, A. Farmer, I. N. Ferrier, E. Flynn, A. Forbes, L. Forty, J. A. Franklyn, T. M. Frayling, R. M. Freathy, E. Giannoulatou, P. Gibbs, P. Gilbert, K. Gordon-Smith, E. Gray, E. Green, C. J. Groves, D. Grozeva, R. Gwilliam, A. Hall, N. Hammond, M. Hardy, P. Harrison, N. Hassanali, H. Hebaishi, S. Hines, A. Hinks, G. A. Hitman, L. Hocking, C. Holmes, E. Howard, P. Howard, J. M. Howson, D. Hughes, S. Hunt, J. D. Isaacs, M. Jain, D. P. Jewell, T. Johnson, J. D. Jolley, I. R. Jones, L. A. Jones, G. Kirov, C. F. Langford, H. Lango-Allen, G. M. Lathrop, J. Lee, K. L. Lee, C. Lees, K. Lewis, C. M. Lindgren, M. Maisuria-Armer, J. Maller, J. Mansfield, J. L. Marchini, P. Martin, D. C. Massey, W. L. McArdle, P. McGuffin, K. E. McLay, G. McVean, A. Mentzer, M. L. Mimmack, A. E. Morgan, A. P. Morris, C. Mowat, P. B. Munroe, S. Myers, W. Newman, E. R. Nimmo, M. C. O'Donovan, A. Onipinla, N. R. Ovington, M. J. Owen, K. Palin, A. Palotie, K. Parnell, R. Pearson, D. Pernet, J. R. Perry, A. Phillips, V. Plagnol, N. J. Prescott, I. Prokopenko, M. A. Quail, S. Rafelt, N. W. Rayner, D. M. Reid, A. Renwick, S. M. Ring, N. Robertson, S. Robson, E. Russell, D. St Clair, J. G. Sambrook, J. D. Sanderson, S. J. Sawcer, H. Schuilenburg, C. E. Scott, R. Scott, S. Seal, S. Shaw-Hawkins, B. M. Shields, M. J. Simmonds, D. J. Smyth, E. Somaskantharajah, K. Spanova, S. Steer, J. Stephens, H. E. Stevens, K. Stirrups, M. A. Stone, D. P. Strachan, Z. Su, D. P. Symmons, J. R. Thompson, W. Thomson, M. D. Tobin, M. E. Travers, C. Turnbull, D. Vukcevic, L. V. Wain, M. Walker, N. M. Walker, C. Wallace, M. Warren-Perry, N. A. Watkins, J. Webster, M. N. Weedon, A. G. Wilson, M. Woodburn, B. P. Wordsworth, C. Yau, A. H. Young, E. Zeggini, M. A. Brown, P. R. Burton, M. J. Caulfield, A. Compston, M. Farrall, S. C. Gough, A. S. Hall, A. T. Hattersley, A. V. Hill, C. G. Mathew, M. Pembrey, J. Satsangi, M. R. Stratton, J. Worthington, M. E. Hurles, A. Duncanson, W. H. Ouwehand, M. Parkes, N. Rahman, J. A. Todd, N. J. Samani, D. P. Kwiatkowski, M. I. McCarthy, N. Craddock, P. Deloukas, P. Donnelly, J. M. Blackwell, E. Bramon, J. P. Casas, A. Corvin, J. Jankowski, H. S. Markus, C. N. Palmer, R. Plomin, A. Rautanen, R. C. Trembath, A. C. Viswanathan, N. W. Wood, C. C. Spencer, G. Band, C. Bellenguez, C. Freeman, G. Hellenthal, E. Giannoulatou, M. Pirinen, R. Pearson, A. Strange, H. Blackburn, S. J. Bumpstead, S. Dronov, M. Gillman, A. Jayakumar, O. T. McCann, J. Liddle, S. C. Potter, R. Ravindrarajah, M. Ricketts, M. Waller, P. Weston, S. Widaa, and P. Whittaker. Host-microbe interactions have shaped the genetic architecture of inflammatory bowel disease. Nature, 491(7422):119–124, Nov 2012.

42 A. Hinks, J. Cobb, M. C. Marion, S. Prahalad, M. Sudman, J. Bowes, P. Martin, M. E. Comeau, S. Sajuthi, R. Andrews, M. Brown, W. M. Chen, P. Concannon, P. Deloukas, S. Edkins, S. Eyre, P. M. Gaffney, S. L. Guthery, J. M. Guthridge, S. E. Hunt, J. A. James, M. Keddache, K. L. Moser, P. A. Nigrovic, S. Onengut-Gumuscu, M. L. Onslow, C. D. Rose, S. S. Rich, K. J. Steel, E. K. Wakeland, C. A. Wallace, L. R. Wedderburn, P. Woo, J. F. Bohnsack, J. P. Haas, D. N. Glass, C. D. Langefeld, W. Thomson, and S. D. Thompson. Dense genotyping of immune-related disease regions identifies 14 new susceptibility loci for juvenile idiopathic arthritis. Nat. Genet., 45(6):664–669, Jun 2013.

43 J. Z. Liu, M. A. Almarri, D. J. Gaffney, G. F. Mells, L. Jostins, H. J. Cordell, S. J. Ducker, D. B. Day, M. A. Heneghan, J. M. Neuberger, P. T. Donaldson, A. J. Bathgate, A. Burroughs, M. H. Davies, D. E. Jones, G. J. Alexander, J. C. Barrett, R. N. Sandford, C. A. Anderson, G. Alexander, A. Bathgate, A. Burroughs, H. Cordell, M. Davies, P. Donaldson, M. Heneghan, D. Jones, G. Mells, J. Neuberger, C. Thain, R. Sandford, B. Street, C. Lye, C. Lai, T. Yapp, R. Sturgess, C. Healey, M. Czajkowski, S. Peter, J. Thornton, S. Mann, K. Kapur, R. Marley, G. Foster, J. Ramage, R. Harvey, N. MacDougall, C. J. Shorrock, G. Lipscomb, P. Southern, N. Parnell, J. Tibble, D. Gorard, G. Mells, M. Dawwas, R. Aspinall, S. Dolwani, M. Foxton, H. Mitchison, I. Gooding, M. Patel, R. Ede, A. Austin, R. Dawood, J. Sayer, C. Hovell, N. Fisher, M. Carter, K. Koss, A. Piotrowicz, D. Banait, D. Neal, G. Lim, A. Ala, A. Saeed, J. Brown, S. Thomas, M. Wilkinson, J. Ridpath, T. Ngatchu, S. Levi, R. Ransford, R. Dickinson, R. Shidrawi, G. Abouda, I. Rees, I. Salam, F. Ali, M. Narain, A. Brown, S. Khakoo, S. Williams, M. Williams, A. Chilton, R. Westbrook, M. Heneghan, C. Rodrigues, M. Davies, M. Aldersley, C. Millson, S. Sen, G. Bird, L. Smith, K. Yoong, N. Rajendran, R. Mathew, G. MacFaul, A. Shah, C. Evans, S. Saha, P. Bramley, A. Fraser, P. Mills, T. Shallcross, D. de Las Heras, C. Sheen, R. Crofton, A. Prach, A. Shepherd, H. Kennedy, S. Rushbrook, R. Przemioslo, C. McDonald, B. Javaid, B. Chaudhury, J. Metcalf, D. Ramanaden, J. Gasem, R. Evans, U. Shmueli, A. Naqvi, J. Collier, H. Klass, M. Ninkovic, M. Cramp, P. Goggin, B. Hoeroldt, G. Lipscomb, E. Williams, H. Hussaini, R. Devon, R. Ayres, J. Makanyanga, A. Burroughs, P. Richardson, M. Lombard, D. Robertson, M. Farrant, A. Tanner, S. Singhal, S. Babu, D. Gleeson, J. Butterworth, K. George, H. Curtis, A. McNair, I. Nasr, A. Douglas, J. Shearman, K. Nash, M. Wright, G. Bray, J. Mclindon, D. Das, G. Whatley, S. Lean, N. Sivaramakrishnan, S. Ducker, D. Jones, D. Preston, A. Douds, M. Brookes, V. S. Wong, S. Pereira, M. Carbone, J. Neuberger, G. Watts, F. Gordon, E. Unitt, A. Grant, M. Cox, S. Whalley, J. Fraser, A. Li, A. Bell, H. Gordon, A. Singhal, I. Ahmad, L. NHS, Y. Ang, J. Gotto, A. Turnbull, C. A. Anderson, J. C. Barrett, J. A. Floyd, C. S, R. McGinnis, N. Soranzo, J. Sambrook, J. Stephens, W. H. Ouwehand, W. L. McArdle, S. M. Ring, D. P. Strachan, G. Alexander, J. C. Barrett, C. M. Bulik, P. J. Conlon, A. Dominiczak, A. Duncanson, A. Hill, G. Lord, A. P. Maxwell, L. Morgan, L. Peltonen, R. N, N. Sheerin, N. Soranzo, F. O. Vannberg, J. C. Barrett, P. Concannon, E. Gray, S. E. Hunt, C. Langford, S. Potter, S. Rich, and D. Simpkin. Dense fine-mapping study identifies new susceptibility loci for primary biliary cirrhosis. Nat. Genet., 44(10):1137–1141, Oct 2012.

44 L. C. Tsoi, S. L. Spain, J. Knight, E. Ellinghaus, P. E. Stuart, F. Capon, J. Ding, Y. Li, T. Tejasvi, J. E. Gudjonsson, H. M. Kang, M. H. Allen, R. McManus, G. Novelli, L. Samuelsson, J. Schalkwijk, M. Stahle, A. D. Burden, C. H. Smith, M. J. Cork, X. Estivill, A. M. Bowcock, G. G. Krueger, W. Weger, J. Worthington, R. Tazi-Ahnini, F. O. Nestle, A. Hayday, P. Hoffmann, J. Winkelmann, C. Wijmenga, C. Langford, S. Edkins, R. Andrews, H. Blackburn, A. Strange, G. Band, R. D. Pearson, D. Vukcevic, C. C. Spencer, P. Deloukas, U. Mrowietz, S. Schreiber, S. Weidinger, S. Koks, K. Kingo, T. Esko, A. Metspalu, H. W. Lim, J. J. Voorhees, M. Weichenthal, H. E. Wichmann, V. Chandran, C. F. Rosen, P. Rahman, D. D. Gladman, C. E. Griffiths, A. Reis, J. Kere, R. P. Nair, A. Franke, J. N. Barker, G. R. Abecasis, J. T. Elder, R. C. Trembath, K. C. Duffin, C. Helms, D. Goldgar, Y. Li, J. Paschall, M. J. Malloy, C. R. Pullinger, J. P. Kane, J. Gardner, A. Perlmutter, A. Miner, B. J. Feng, R. Hiremagalore, R. W. Ike, E. Christophers, T. Henseler, A. Ruether, S. J. Schrodi, S. Prahalad, S. L. Guthery, J. Fischer, W. Liao, P. Kwok, A. Menter, G. M. Lathrop, C. Wise, A. B. Begovich, A. Onoufriadis, M. E. Weale, A. Hofer, W. Salmhofer, P. Wolf, K. Kainu, U. Saarialho-Kere, S. Suomela, P. Badorf, U. Huffmeier, W. Kurrat, W. Kuster, J. Lascorz, R. Mossner, F. Schurmeier-Horst, M. Stander, H. Traupe, J. G. Bergboer, M. den Heijer, P. C. van de Kerkhof, P. L. Zeeuwen, L. Barnes, L. E. Campbell, C. Cusack, C. Coleman, J. Conroy, S. Ennis, O. Fitzgerald, P. Gallagher, A. D. Irvine, B. Kirby, T. Markham, W. H. McLean, J. McPartlin, S. F. Rogers, A. W. Ryan, A. Zawirska, E. Giardina, T. Lepre, C. Perricone, G. Martin-Ezquerra, R. M. Pujol, E. Riveira-Munoz, A. Inerot, A. T. Naluai, L. Mallbris, K. Wolk, J. Leman, A. Barton, R. B. Warren, H. S. Young, I. Ricano-Ponce, G. Trynka, F. J. Pellett, A. Henschel, M. Aurand, B. Bebo, C. Gieger, T. Illig, S. Moebus, K. H. Jockel, R. Erbel, P. Donnelly, L. Peltonen, J. M. Blackwell, E. Bramon, M. A. Brown, J. P. Casas, A. Corvin, N. Craddock, A. Duncanson, J. Jankowski, H. S. Markus, C. G. Mathew, M. I. McCarthy, C. N. Palmer, R. Plomin, A. Rautanen, S. J. Sawcer, N. Samani, A. C. Viswanathan, N. W. Wood, C. Bellenguez, C. Freeman, G. Hellenthal, E. Giannoulatou, M. Pirinen, Z. Su, S. E. Hunt, R. Gwilliam, S. J. Bumpstead, S. Dronov, M. Gillman, E. Gray, N. Hammond, A. Jayakumar, O. T. McCann, J. Liddle, M. L. Perez, S. C. Potter, R. Ravindrarajah, M. Ricketts, M. Waller, P. Weston, S. Widaa, and P. Whittaker. Identification of 15 new psoriasis susceptibility loci highlights the role of innate immunity. Nat. Genet., 44(12):1341–1348, Dec 2012.

45 Y. Okada, D. Wu, G. Trynka, T. Raj, C. Terao, K. Ikari, Y. Kochi, K. Ohmura, A. Suzuki, S. Yoshida, R. R. Graham, A. Manoharan, W. Ortmann, T. Bhangale, J. C. Denny, R. J. Carroll, A. E. Eyler, J. D. Greenberg, J. M. Kremer, D. A. Pappas, L. Jiang, J. Yin, L. Ye, D. F. Su, J. Yang, G. Xie, E. Keystone, H. J. Westra, T. Esko, A. Metspalu, X. Zhou, N. Gupta, D. Mirel, E. A. Stahl, D. Diogo, J. Cui, K. Liao, M. H. Guo, K. Myouzen, T. Kawaguchi, M. J. Coenen, P. L. van Riel, M. A. van de Laar, H. J. Guchelaar, T. W. Huizinga, P. Dieude, X. Mariette, S. L. Bridges, A. Zhernakova, R. E. Toes, P. P. Tak, C. Miceli-Richard, S. Y. Bang, H. S. Lee, J. Martin, M. A. Gonzalez-Gay, L. Rodriguez-Rodriguez, S. Rantapaa-Dahlqvist, L. Arlestig, H. K. Choi, Y. Kamatani, P. Galan, M. Lathrop, S. Eyre, J. Bowes, A. Barton, N. de Vries, L. W. Moreland, L. A. Criswell, E. W. Karlson, A. Taniguchi, R. Yamada, M. Kubo, J. S. Liu, S. C. Bae, J. Worthington, L. Padyukov, L. Klareskog, P. K. Gregersen, S. Raychaudhuri, B. E. Stranger, P. L. De Jager, L. Franke, P. M. Visscher, M. A. Brown, H. Yamanaka, T. Mimori, A. Takahashi, H. Xu, T. W. Behrens, K. A. Siminovitch, S. Momohara, F. Matsuda, K. Yamamoto, R. M. Plenge, S. Eyre, J. Bowes, D. Diogo, A. Lee, A. Barton, P. Martin, A. Zhernakova, E. Stahl, S. Viatte, K. McAllister, C. I. Amos, L. Padyukov, R. E. Toes, T. W. Huizinga, C. Wijmenga, G. Trynka, L. Franke, H. J. Westra, L. Alfredsson, X. Hu, C. Sandor, P. I. de Bakker, S. Davila, C. C. Khor, K. K. Heng, R. Andrews, S. Edkins, S. E. Hunt, C. Langford, D. Symmons, P. Concannon, S. Onengut-Gumuscu, S. S. Rich, P. Deloukas, M. A. Gonzalez-Gay, L. Rodriguez-Rodriguez, L. Arlsetig, J. Martin, S. Rantapaa-Dahlqvist, R. M. Plenge, S. Raychaudhuri, L. Klareskog, P. K. Gregersen, J. Worthington, Y. Okada, C. Terao, K. Ikari, Y. Kochi, K. Ohmura, A. Suzuki, T. Kawaguchi, E. Stahl, F. Kurreman, N. Nishida, H. Ohmiya, K. Myouzen, M. Takahashi, T. Sawada, Y. Nishioka, M. Yukioka, T. Matsubara, S. Wakitani, R. Teshima, S. Tohma, K. Takasugi, K. Shimada, A. Murasawa, S. Honjo, K. Matsuo, H. Tanaka, K. Tajima, T. Suzuki, T. Iwamoto, Y. Kawamura, H. Tanii, Y. Okazaki, T. Sasaki, P. K. Gregersen, L. Padyukov, J. Worthington, K. A. Siminovitch, M. Lathrop, A. Taniguchi, A. Takahashi, K. Tokunaga, M. Kubo, Y. Nakamura, N. Kamatani, T. Mimori, R. M. Plenge, H. Yamanaka, S. Momohara, R. Yamada, F. Matsuda, and K. Yamamoto. Genetics of rheumatoid arthritis contributes to biology and drug discovery. Nature, 506(7488):376–381, Feb 2014.

46 S. Onengut-Gumuscu, W. M. Chen, O. Burren, N. J. Cooper, A. R. Quinlan, J. C. Mychaleckyj, E. Farber, J. K. Bonnie, M. Szpak, E. Schofield, P. Achuthan, H. Guo, M. D. Fortune, H. Stevens, N. M. Walker, L. D. Ward, A. Kundaje, M. Kellis, M. J. Daly, J. C. Barrett, J. D. Cooper, P. Deloukas, J. A. Todd, C. Wallace, P. Concannon, S. S. Rich, T. Baskerville, N. Bautista, E. Bhatia, V. Bhatia, K. Bin Hasan, F. Bonnici, T. Brodnicki, B. Browning, F. Cameron, K. Chaichanwatanakul, P. T. Cheung, P. Colman, A. Cotterill, J. Couper, P. Crock, R. Cutfield, T. Davis, P. Dixon, K. Donaghue, K. Dowling, P. Drury, S. Dye, S. Gellert, R. A. Ghani, R. Greer, X. Han, L. Harrison, N. Homatopoulos, L. Ji, T. Jones, L. K. Yin, N. A. Kamaruddin, U. Kanga, A. Kanungo, G. Kaur, B. Kek, S. Knowles, J. Krebs, N. Kumar, Y. J. Lee, X. Li, S. Liktimaskul, M. Lloyd, A. Loth, A. Louey, N. Mehra, T. Merriman, L. Min, G. Morahan, R. Moses, G. Mraz, R. Murphy, I. Nicholson, A. Panelo, P. Poh, G. Price, N. Ratnam, C. Sanjeevi, S. Sedimbi, S. Shen, G. Siok Ying, B. Tait, N. Tandon, A. Thomas, M. Varney, P. Weerakulwattana, J. Willis, E. A. Akwo, L. Albret, F. Ampudia-Blasco, J. Argente, M. Avbelj, G. Babadjanova, K. Badenhoop, T. Battelino, G. Beilhack, R. Bergholdt, P. Bingley, B. Boehm, J. Bolidson, K. Brismar, C. Brorsson, J. Carlson, L. Castano, K. Chandler, V. Cherubini, O. Cinek, E. Cipponeri, R. Corripio Collado, A. de Leiva, I. Dzivite, A. Fagulha, M. Fernandez Balcells, B. Garcia Cuartero, C. Garcia Lacalle, C. Guja, P. Gutierrez, A. Hamou, E. Hatziagelaki, S. Heath, K. Heilman, W. Helmberg, O. Hermon, M. Hernandez, I. Holzheu, N. Hosszufalusi, J. Ilonen, C. Ionescu-Tirgoviste, J. Johannesen, C. Julier, H. Kahles, I. Kinalska, M. Knip, I. Kockum, E. Kojo, O. Kordonouri, A. Kretowski, D. Krikovszky, A. Kurkhaus, M. Kuzmicki, E. Lavant, A. Long, J. Ludvigsson, L. Madacsy, K. Maliszewska, M. Marga, M. P. Martinez, D. Mauricio, G. Mazurkievicz, J. Nerup, A. Norkus, F. J. Novoa Mogollon, A. Okruszko, C. Pettinari, M. Phillip, V. Pirags, F. Pociot, P. Pozzilli, R. Racasan, K. Raile, R. Rappner, M. J. Rodriguez Troyano, B. O. Roep, S. Rokni, S. Rosinger, O. Rubio-Cabezas, C. Ruckgaber, I. Satman, E. Schober, J. Seufert, R. Sing, J. Skrha, E. Sobngwi, M. Somerville, G. Spinas, Z. Sumnik, V. Tilmann, D. Undlien, V. Urbanavicius, B. Van der Auwera, F. Vasquez San Miguel, A. Vazeo-Gerasimidi, D. Velickiene, A. Wagner, M. Walter, A. Williams, A. Ziegler, M. Agleham, A. Aldrich, R. Alemzadeh, C. Alper, T. Aly, D. Anastassiou, S. Arora, A. Austin, D. Becker, C. Benoist, N. Berka, S. Bhatia, P. Bonella, N. Bottini, S. Boyle, J. Braden, B. Brady, W. Brickman, R. Christensen, P. Concannon, R. Couch, D. Counts, J. Crandall, M. Daniels, L. Dolan, D. Donaldson, A. Doria, G. Eisenbarth, J. Elder, R. El-Hajj, H. Erlich, P. Fain, A. L. Fear, R. Ferry, R. Fiallo-Scharer, D. Geraghty, S. Ghosh, S. Gitelman, M. Godwin, R. Goland, N. Goodman, G. Goodwin, J. Gravely, C. Greenbaum, C. Gudgeon, F. Gunville, W. Hagopian, H. Hakonarson, J. Hansen, K. Harrington, J. Hassing, W. Hilliker, R. Hoffman, E. Hulbert, R. Izquierdo, N. Jospe, K. Kaiserman, F. Kaufman, S. Kim, E. Kloos, R. Kosoy, J. Lane, J. Lane, J. Lawrence, C. Levetan, P. Levin, R. Lipton, J. Lonsdale, V. Magnuson, J. Marks, B. Mayer-Davis, R. McEvoy, R. McIndoe, L. Merkle, D. Metzger, D. Miao, E. Mickelson, P. Moonsamy, W. Moore, A. Moran, J. Noble, G. Olsem, S. Onengut-Gumuscu, T. Orban, C. Orlowski, A. Paterson, M. Pietropaolo, C. Pihoker, C. Polychronakos, J. Post, D. Postellon, A. Pugliese, H. Qu, T. Quattrin, M. Rappaport, P. Raskin, H. Risbeck, H. Rodriguez, L. Rodriguez, M. Rogers, L. Rubalcava, B. Russell, D. Schatz, C. Scott, J. X. She, H. Shilling, D. Shulman, L. Soyka, P. Speiser, H. Starkman, A. Steck, S. Stender, L. Stratton, D. Sur, S. Taback, K. Thrailkill, E. Toth, P. Trymbiski, E. Tsalikian, K. Vertachnik, J. Wahlen, X. Wang, S. Weber, D. Wherrett, S. Willi, D. Wilson, J. Youkey, N. Young, L. Yu, L. P. Zhao, D. Zimmerman, E. Adlem, J. Allen, J. Barrett, J. Brown, O. Burren, P. Clarke, D. Clayton, G. Coleman, J. Cooper, F. Cucca, L. Davison, K. Downes, S. Duley, D. Dunger, L. Esposito, V. Everett, S. Field, J. Hafler, M. Hardy, D. Harrison, I. Harrison, S. Hawkins, B. Healy, S. Hood, S. Howell, J. Howson, M. Maisuria, W. Meadows, T. Mistry, S. Nezhenstsev, S. Nutland, N. Ovington, V. Plagnol, D. Rainbow, K. Rainbow, S. Raj, H. Schuilenburg, A. Simpson, L. Smink, D. Smyth, H. Stevens, N. Taylor, J. Todd, J. Tuomilehto, N. Walker, L. Wicker, B. Widmer, M. Wilson, H. Withers, J. Yang, M. Brown, W. M. Chen, A. Crews, J. Griffin, M. Hall, T. Harnish, J. Hepler, J. Hilner, N. King, K. Lohman, L. Lu, J. Mychaleckyj, J. Nail, L. Perdue, J. Pierce, D. Reboussin, S. Rich, S. Rushing, M. Sale, E. Sides, B. Snively, H. Teuschler, G. Theil, L. Wagenknecht, D. Williams, B. Akolkar, C. McKeon, C. Nierras, E. Thomson, D. Altshuler, K. Au, S. Bain, L. Barcellos, S. Barral, T. Becker, F. Briggs, P. Bronson, M. Daly, P. de Bakker, P. Deloukas, B. Devlin, M. C. Eike, L. Field, S. Gabriel, N. Garge, S. Gaudieri, B. Goldstein, C. Gorodezky, S. Hamon, C. He, J. Howson, K. Humphreys, I. James, M. Lathrop, B. A. Lie, D. Li, S. Mack, R. McGinnis, E. McKinnon, W. McLaren, D. Nolan, M. Olsson, J. Ott, D. Owerbach, C. Patterson, R. Podolsky, P. Ramsay, V. Rangantah, N. Risch, K. S. Ronningen, X. Shao, R. Single, M. Steffes, G. Thomson, A. M. Valdes, C. Vandiedonck, P. Whittaker, and Q. Zhang. Fine mapping of type 1 diabetes susceptibility loci and evidence for colocalization of causal variants with lymphoid gene enhancers. Nat. Genet., 47(4):381–386, Apr 2015.

47 H. J. Cordell, Y. Han, G. F. Mells, Y. Li, G. M. Hirschfield, C. S. Greene, G. Xie, B. D. Juran, D. Zhu, D. C. Qian, J. A. Floyd, K. I. Morley, D. Prati, A. Lleo, D. Cusi, M. E. Gershwin, C. A. Anderson, K. N. Lazaridis, P. Invernizzi, M. F. Seldin, R. N. Sandford, C. I. Amos, K. A. Siminovitch, E. M. Schlicht, C. Lammert, E. J. Atkinson, L. L. Chan, M. de Andrade, T. Balschun, A. L. Mason, R. P. Myers, J. Zhang, P. Milkiewicz, J. Qu, J. A. Odin, V. A. Luketic, B. R. Bacon, H. C. Bodenheimer, V. Liakina, C. Vincent, C. Levy, P. K. Gregersen, P. L. Almasio, D. Alvaro, P. Andreone, A. Andriulli, C. Barlassina, P. M. Battezzati, A. Benedetti, F. Bernuzzi, I. Bianchi, M. C. Bragazzi, M. Brunetto, S. Bruno, G. Casella, B. Coco, A. Colli, M. Colombo, S. Colombo, C. Cursaro, L. S. Croce, A. Crosignani, M. F. Donato, G. Elia, L. Fabris, C. Ferrari, A. Floreani, B. Foglieni, R. Fontana, A. Galli, R. Lazzari, F. Macaluso, F. Malinverno, F. Marra, M. Marzioni, A. Mattalia, R. Montanari, L. Morini, F. Morisco, M. Hani S, L. Muratori, P. Muratori, G. A. Niro, V. O. Palmieri, A. Picciotto, M. Podda, P. Portincasa, V. Ronca, F. Rosina, S. Rossi, I. Sogno, G. Spinzi, M. Spreafico, M. Strazzabosco, S. Tarallo, M. Tarocchi, C. Tiribelli, P. Toniutto, M. Vinci, M. Zuin, C. L. Ch'ng, M. Rahman, T. Yapp, R. Sturgess, C. Healey, M. Czajkowski, A. Gu-nasekera, P. Gyawali, P. Premchand, K. Kapur, R. Marley, G. Foster, A. Watson, A. Dias, J. Subhani, R. Harvey, R. McCorry, D. Ramanaden, J. Gasem, R. Evans, T. Mathialahan, C. Shorrock, G. Lipscomb, P. Southern, J. Tibble, D. Gorard, A. Palegwala, S. Jones, M. Carbone, M. Dawwas, G. Alexander, S. Dolwani, M. Prince, M. Foxton, D. Elphick, H. Mitchison, I. Gooding, M. Karmo, S. Saksena, M. Mendall, M. Patel, R. Ede, A. Austin, J. Sayer, L. Hankey, C. Hovell, N. Fisher, M. Carter, K. Koss, A. Piotrowicz, C. Grimley, D. Neal, G. Lim, S. Levi, A. Ala, A. Broad, A. Saeed, G. Wood, J. Brown, M. Wilkinson, H. Gordon, J. Ramage, J. Ridpath, T. Ngatchu, B. Grover, S. Shaukat, R. Shidrawi, G. Abouda, F. Ali, I. Rees, I. Salam, M. Narain, A. Brown, S. Taylor-Robinson, S. Williams, L. Grellier, P. Banim, D. Das, A. Chilton, M. Heneghan, H. Curtis, M. Gess, I. Drake, M. Aldersley, M. Davies, R. Jones, A. McNair, R. Srirajaskanthan, M. Pitcher, S. Sen, G. Bird, A. Barnardo, P. Kitchen, K. Yoong, J. Chirag, N. Sivaramakrishnan, G. MacFaul, D. Jones, A. Shah, C. Evans, S. Saha, K. Pollock, P. Bramley, A. Mukhopadhya, A. Fraser, P. Mills, C. Shallcross, S. Campbell, A. Bathgate, A. Shepherd, J. Dillon, S. Rushbrook, R. Przemioslo, C. Macdonald, J. Metcalf, U. Shmueli, A. Davis, A. Naqvi, T. Lee, S. D. Ryder, J. Collier, H. Klass, M. Ninkovic, M. Cramp, N. Sharer, R. Aspinall, P. Goggin, D. Ghosh, A. Douds, B. Hoeroldt, J. Booth, E. Williams, H. Hussaini, W. Stableforth, R. Ayres, D. Thorburn, E. Marshall, A. Burroughs, S. Mann, M. Lombard, P. Richardson, I. Patanwala, J. Maltby, M. Brookes, R. Mathew, S. Vyas, S. Singhal, D. Gleeson, S. Misra, J. Butterworth, K. George, T. Harding, A. Douglass, S. Panter, J. Shearman, G. Bray, G. Butcher, D. Forton, J. Mclindon, M. Cowan, G. Whatley, A. Mandal, H. Gupta, P. Sanghi, S. Jain, S. Pereira, G. Prasad, G. Watts, M. Wright, J. Neuberger, F. Gordon, E. Unitt, A. Grant, T. Delahooke, A. Higham, A. Brind, M. Cox, S. Ramakrishnan, A. King, C. Collins, S. Whalley, A. Li, J. Fraser, A. Bell, V. S. Wong, A. Singhal, I. Gee, Y. Ang, R. Ransford, J. Gotto, C. Millson, J. Bowles, C. Thomas, M. Harrison, R. Galaska, J. Kendall, J. Whiteman, C. Lawlor, C. Gray, K. Elliott, C. Mulvaney-Jones, L. Hobson, G. Van Duyvenvoorde, A. Loftus, K. Seward, R. Penn, J. Maiden, R. Damant, J. Hails, R. Cloudsdale, V. Silvestre, S. Glenn, E. Dungca, N. Wheatley, H. Doyle, M. Kent, C. Hamilton, D. Braim, H. Wooldridge, R. Abrahams, A. Paton, N. Lancaster, A. Gibbins, K. Hogben, P. Desousa, F. Muscariu, J. Musselwhite, A. McKay, L. Tan, C. Foale, J. Brighton, K. Flahive, E. Nambela, P. Townshend, C. Ford, S. Holder, C. Palmer, J. Featherstone, M. Nasseri, J. Sadeghian, B. Williams, C. Thomas, S. A. Rolls, A. Hynes, C. Duggan, S. Jones, M. Crossey, G. Stansfield, C. MacNicol, J. Wilkins, E. Wilhelmsen, P. Raymode, H. J. Lee, E. Durant, R. Bishop, N. Ncube, S. Tripoli, R. Casey, C. Cowley, R. Miller, K. Houghton, S. Ducker, F. Wright, B. Bird, G. Baxter, J. Keggans, M. Hughes, E. Grieve, K. Young, D. Williams, K. Ocker, F. Hines, K. Martin, C. Innes, T. Valliani, H. Fairlamb, S. Thornthwaite, A. Eastick, E. Tanqueray, J. Morrison, B. Holbrook, J. Browning, K. Walker, S. Congreave, J. Verheyden, S. Slininger, L. Stafford, D. O'Donnell, M. Ainsworth, S. Lord, L. Kent, L. March, C. Dickson, D. Simpson, B. Longhurst, M. Hayes, E. Shpuza, N. White, S. Besley, S. Pearson, A. Wright, L. Jones, E. Gunter, H. Dewhurst, A. Fouracres, L. Farrington, L. Graves, S. Marriott, M. Leoni, D. Tyrer, K. Martin, L. Dali-Kemmery, V. Lambourne, M. Green, D. Sirdefield, K. Amor, J. Colley, B. Shinder, J. Jones, M. Mills, M. Carnahan, N. Taylor, K. Boulton, J. Tregonning, C. Brown, G. Clifford, E. Archer, M. Hamilton, J. Curtis, T. Shewan, S. Walsh, K. Warner, K. Netherton, M. Mupudzi, B. Gunson, J. Gitahi, D. Gocher, S. Batham, H. Pateman, S. Desmennu, J. Conder, D. Clement, S. Gallagher, J. Orpe, P. Chan, L. Currie, L. O'Donohoe, M. Oblak, L. Morgan, M. Quinn,I. Amey, Y. Baird, D. Cotterill, L. Cumlat, L. Winter, S. Greer, K. Spurdle, J. Allison, S. Dyer, H. Sweeting, and J. Kordula. International genome-wide meta-analysis identifies new primary biliary cirrhosis risk loci and targetable pathogenic pathways. Nat Commun, 6:8019, 2015.

48 G. R. Abecasis, D. Altshuler, A. Auton, L. D. Brooks, R. M. Durbin, R. A. Gibbs, M. E. Hurles, G. A. McVean, D. Altshuler, R. M. Durbin, G. R. Abecasis, D. R. Bentley, A. Chakravarti, A. G. Clark, F. S. Collins, F. M. De La Vega, P. Donnelly, M. Egholm, P. Flicek, S. B. Gabriel, R. A. Gibbs, B. M. Knoppers, E. S. Lander, H. Lehrach, E. R. Mardis, G. A. McVean, D. A. Nickerson, L. Peltonen, A. J. Schafer, S. T. Sherry, J. Wang, R. Wilson, R. A. Gibbs, D. Deiros, M. Metzker, D. Muzny, J. Reid, D. Wheeler, J. Wang, J. Li, M. Jian, G. Li, R. Li, H. Liang, G. Tian, B. Wang, J. Wang, W. Wang, H. Yang, X. Zhang, H. Zheng, E. S. Lander, D. L. Altshuler, L. Ambrogio, T. Bloom, K. Cibulskis, T. J. Fennell, S. B. Gabriel, D. B. Jaffe, E. Shefler, C. L. Sougnez, D. R. Bentley, N. Gormley, S. Humphray, Z. Kingsbury, P. Kokko-Gonzales, J. Stone, K. J. McKernan, G. L. Costa, J. K. Ichikawa, C. C. Lee, R. Sudbrak, H. Lehrach, T. A. Borodina, A. Dahl, A. N. Davydov, P. Marquardt, F. Mertes, W. Nietfeld, P. Rosenstiel, S. Schreiber, A. V. Soldatov, B. Timmermann, M. Tolzmann, M. Egholm, J. Affourtit, D. Ashworth, S. Attiya, M. Bachorski, E. Buglione, A. Burke, A. Caprio, C. Celone, S. Clark, D. Conners, B. Desany, L. Gu, L. Guccione, K. Kao, J. Kebbler, J. Knowlton, M. Labrecque, L. McDade, C. Mealmaker, M. Minderman, A. Nawrocki, F. Niazi, K. Pareja, R. Ramenani, D. Riches, W. Song, C. Turcotte, S. Wang, E. R. Mardis, R. K. Wilson, D. Dooling, L. Fulton, R. Fulton, G. Weinstock, R. M. Durbin, J. Burton, D. M. Carter, C. Churcher, A. Coffey, A. Cox, A. Palotie, M. Quail, T. Skelly, J. Stalker, H. P. Swerdlow, D. Turner, A. De Witte, S. Giles, R. A. Gibbs, D. Wheeler, M. Bainbridge, D. Challis, A. Sabo, F. Yu, J. Yu, J. Wang, X. Fang, X. Guo, R. Li, Y. Li, R. Luo, S. Tai, H. Wu, H. Zheng, X. Zheng, Y. Zhou, G. Li, J. Wang, H. Yang, G. T. Marth, E. P. Garrison, W. Huang, A. Indap, D. Kural, W. P. Lee, W. F. Leong, A. R. Quinlan, C. Stewart, M. P. Stromberg, A. N. Ward, J. Wu, C. Lee, R. E. Mills, X. Shi, M. J. Daly, M. A. DePristo, D. L. Altshuler, A. D. Ball, E. Banks, T. Bloom, B. L. Browning, K. Cibulskis, T. J. Fennell, K. V. Garimella, S. R. Grossman, R. E. Handsaker, M. Hanna, C. Hartl, D. B. Jaffe, A. M. Kernytsky, J. M. Korn, H. Li, J. R. Maguire, S. A. McCarroll, A. McKenna, J. C. Nemesh, A. A. Philippakis, R. E. Poplin, A. Price, M. A. Rivas, P. C. Sabeti, S. F. Schaffner, E. Shefler, I. A. Shlyakhter, D. N. Cooper, E. V. Ball, M. Mort, A. D. Phillips, P. D. Stenson, J. Sebat, V. Makarov, K. Ye, S. C. Yoon, C. D. Bustamante, A. G. Clark, A. Boyko, J. Degenhardt, S. Gravel, R. N. Gutenkunst, M. Kaganovich, A. Keinan, P. Lacroute, X. Ma, A. Reynolds, L. Clarke, P. Flicek, F. Cunningham, J. Herrero, S. Keenen, E. Kulesha, R. Leinonen, W. M. McLaren, R. Radhakrishnan, R. E. Smith, V. Zalunin, X. Zheng-Bradley, J. O. Korbel, A. M. Stutz, S. Humphray, M. Bauer, R. K. Cheetham, T. Cox, M. Eberle, T. James, S. Kahn, L. Murray, A. Chakravarti, K. Ye, F. M. De La Vega, Y. Fu, F. C. Hyland, J. M. Manning, S. F. McLaughlin, H. E. Peckham, O. Sakarya, Y. A. Sun, E. F. Tsung, M. A. Batzer, M. K. Konkel, J. A. Walker, R. Sudbrak, M. W. Albrecht, V. S. Amstislavskiy, R. Herwig, D. V. Parkhomchuk, S. T. Sherry, R. Agarwala, H. M. Khouri, A. O. Morgulis, J. E. Paschall, L. D. Phan, K. E. Rotmistrovsky, R. D. Sanders, M. F. Shumway, C. Xiao, G. A. McVean, A. Auton, Z. Iqbal, G. Lunter, J. L. Marchini, L. Moutsianas, S. Myers, A. Tumian, B. Desany, J. Knight, R. Winer, D. W. Craig, S. M. Beckstrom-Sternberg, A. Christoforides, A. A. Kurdoglu, J. V. Pearson, S. A. Sinari, W. D. Tembe, D. Haussler, A. S. Hinrichs, S. J. Katzman, A. Kern, R. M. Kuhn, M. Przeworski, R. D. Hernandez, B. Howie, J. L. Kelley, S. C. Melton, G. R. Abecasis, Y. Li, P. Anderson, T. Blackwell, W. Chen, W. O. Cookson, J. Ding, H. M. Kang, M. Lathrop, L. Liang, M. F. Moffatt, P. Scheet, C. Sidore, M. Snyder, X. Zhan, S. Zollner, P. Awadalla, F. Casals, Y. Idaghdour, J. Keebler, E. A. Stone, M. Zilversmit, L. Jorde, J. Xing, E. E. Eichler, G. Aksay, C. Alkan, I. Hajirasouliha, F. Hormozdiari, J. M. Kidd, S. C. Sahinalp, P. H. Sudmant, E. R. Mardis, K. Chen, A. Chinwalla, L. Ding, D. C. Koboldt, M. D. McLellan, D. Dooling, G. Weinstock, J. W. Wallis, M. C. Wendl, Q. Zhang, R. M. Durbin, C. A. Albers, Q. Ayub, S. Balasubramaniam, J. C. Barrett, D. M.Carter, Y. Chen, D. F. Conrad, P. Danecek, E. T. Dermitzakis, M. Hu, N. Huang, M. E. Hurles, H. Jin, L. Jostins, T. M. Keane, S. Q. Le, S. Lindsay, Q. Long, D. G. MacArthur, S. B. Montgomery, L. Parts, J. Stalker, C. Tyler-Smith, K. Walter, Y. Zhang, M. B. Gerstein, M. Snyder, A. Abyzov, S. Balasubramanian, R. Bjornson, J. Du, F. Grubert, L. Habegger, R. Haraksingh, J. Jee, E. Khurana, H. Y. Lam, J. Leng, X. J. Mu, A. E. Urban, Z. Zhang, Y. Li, R. Luo, G. T. Marth, E. P. Garrison, D. Kural, A. R. Quinlan, C. Stewart, M. P. Stromberg, A. N. Ward, J. Wu, C. Lee, R. E. Mills, X. Shi, S. A. McCarroll, E. Banks, M. A. DePristo, R. E. Handsaker, C. Hartl, J. M. Korn, H. Li, J. C. Nemesh, J. Sebat, V. Makarov, K. Ye, S. C. Yoon, J. Degenhardt, M. Kaganovich, L. Clarke, R. E. Smith, X. Zheng-Bradley, J. O. Korbel, S. Humphray, R. K. Cheetham, M. Eberle, S. Kahn, L. Murray, K. Ye, F. M. De La Vega, Y. Fu, H. E. Peckham, Y. A. Sun, M. A. Batzer, M. K. Konkel, J. A. Walker, C. Xiao, Z. Iqbal, B. Desany, T. Blackwell, M. Snyder, J. Xing, E. E. Eichler, G. Aksay, C. Alkan, I. Hajirasouliha, F. Hormozdiari, J. M. Kidd, K. Chen, A. Chinwalla, L. Ding, M. D. McLellan, J. W. Wallis, M. E. Hurles, D. F. Conrad, K. Walter, Y. Zhang, M. B. Gerstein, M. Snyder, A. Abyzov, J. Du, F. Grubert, R. Haraksingh, J. Jee, E. Khurana, H. Y. Lam, J. Leng, X. J. Mu, A. E. Urban, Z. Zhang, R. A. Gibbs, M. Bainbridge, D. Challis, C. Coafra, H. Dinh, C. Kovar, S. Lee, D. Muzny, L. Nazareth, J. Reid, A. Sabo, F. Yu, J. Yu, G. T. Marth, E. P. Garrison, A. Indap, W. F. Leong, A. R. Quinlan, C. Stewart, A. N. Ward, J. Wu, K. Cibulskis, T. J. Fennell, S. B. Gabriel, K. V. Garimella, C. Hartl, E. Shefler, C. L. Sougnez, J. Wilkinson, A. G. Clark, S. Gravel, F. Grubert, L. Clarke, P. Flicek, R. E. Smith, X. Zheng-Bradley, S. T. Sherry, H. M. Khouri, J. E. Paschall, M. F. Shumway, C. Xiao, G. A. McVean, S. J. Katzman, G. R. Abecasis, E. R. Mardis, D. Dooling, L. Fulton, R. Fulton, D. C. Koboldt, R. M. Durbin, S. Balasubramaniam, A. Coffey, T. M. Keane, D. G. MacArthur, A. Palotie, C. Scott, J. Stalker, C. Tyler-Smith, M. B. Gerstein, S. Balasubramanian, A. Chakravarti, B. M. Knoppers, G. R. Abecasis, C. D. Bustamante, N. Gharani, R. A. Gibbs, L. Jorde, J. S. Kaye, A. Kent, T. Li, A. L. McGuire, G. A. McVean, P. N. Ossorio, C. N. Rotimi, Y. Su, L. H. Toji, C. Tyler-Smith, L. D. Brooks, A. L. Felsenfeld, J. E. McEwen, A. Abdallah, C. R. Juenger, N. C. Clemm, S. Collins, A. Duncanson, E. D. Green, M. S. Guyer, J. L. Peterson, A. J. Schafer, Y. Xue, and R. A. Cartwright. A map of human genome variation from population-scale sequencing. Nature, 467(7319):1061–1073, Oct 2010.

